# Enhanced Mirror Neuron Network Activity and Effective Connectivity during Live Interaction Among Female Subjects

**DOI:** 10.1101/2022.04.22.489113

**Authors:** Chun-Ting Hsu, Wataru Sato, Takanori Kochiyama, Ryusuke Nakai, Kohei Asano, Nobuhito Abe, Sakiko Yoshikawa

## Abstract

Facial expressions are indispensable in daily human communication. Previous neuroimaging studies investigating facial expression processing have presented pre-recorded stimuli and lacked live face-to-face interaction. Our paradigm alternated between presentations of real-time model performance and pre-recorded videos of dynamic facial expressions to participants. Simultaneous functional magnetic resonance imaging (fMRI) and facial electromyography activity recordings, as well as post-scan valence and arousal ratings were acquired from 44 female participants. Live facial expressions enhanced the subjective valence and arousal ratings as well as facial muscular responses. Live performances showed greater engagement of the right posterior superior temporal sulcus (pSTS), right inferior frontal gyrus (IFG), right amygdala and right fusiform gyrus, and modulated the effective connectivity within the right mirror neuron system (IFG, pSTS, and right inferior parietal lobule). A support vector machine algorithm could classify multivoxel activation patterns in brain regions involved in dynamic facial expression processing in the mentalizing networks (anterior and posterior cingulate cortex). These results indicate that live social interaction modulates the activity and connectivity of the right mirror neuron system and enhances spontaneous mimicry, further facilitating emotional contagion.

**Highlights:**

- We alternately presented real-time and pre-recorded dynamic facial expressions.
- Live facial expressions enhanced emotion contagion and spontaneous facial mimicry.
- Live conditions modulated mirror neuron system activity and effective connectivity.
- The mentalizing network showed distinctive multivoxel patterns in live conditions.
- The results support the validity of second-person design in social neuroscience.

## 1. Introduction

Facial expressions are an integral component of emotional communication and interaction (Adolphs, 2002). They direct the receiver’s attention toward, and evoke inferences about, the sender’s affective state and expression in communication (Fernández-Dols, 2017). Facial expression observation evokes spontaneous facial mimicry (Dimberg, 1982), which, in the affiliative social context (Hess, 2021; Hess and Fischer, 2013), facilitates emotional contagion and can act as an implicit expression of affective empathy (Arnold and Winkielman, 2020; Gonzalez-Liencres et al., 2013; Herrando and Constantinides, 2021; Prochazkova and Kret, 2017). Both motor synchrony and emotional contagion contribute to social alignment and herding behaviors (Shamay-Tsoory et al., 2019). The extent of spontaneous facial mimicry and emotional contagion, as estimated in subjective affective experiences, is associated with the liveliness of the stimuli. Pre-recorded dynamic emotional facial expressions have been shown to elicit stronger subjective arousal and more evident facial mimicry than static pictures of emotional facial expressions (Sato and Yoshikawa, 2007a, 2007b). In comparison with viewing pre-recorded dynamic facial expressions, the observation of dynamic facial expressions in live interactions further potentiates subjective emotional experience in both valence and arousal dimensions, as well as the amplitude of spontaneous facial mimicry (Hietanen et al., 2019; Hsu et al., 2020).

Previous neuroimaging studies investigating the processing of dynamic facial expressions have identified the involvement of several neural substrates, such as the superior temporal sulcus (STS) (Allison et al., 2000), as well as the adjacent superior and middle temporal gyri, fusiform gyrus, inferior frontal gyrus (IFG) including premotor areas, and limbic regions including the amygdala (Zinchenko et al., 2018). These brain regions belong to three neural networks that are particularly relevant to social interaction: the mirror neuron system (MNS), mentalizing/theory of mind (ToM) network, and salience network. The MNS, which involves the IFG, ventral premotor cortex, inferior parietal lobe (IPL), and pSTS, is involved in social motor information processing and supports mimicry behaviors (Hamilton, 2008; Iacoboni and Dapretto, 2006; Molenberghs et al., 2012, 2009). To explain the MNS involvement in mimicry, Hamilton proposed that the pSTS processes the kinematic features of observed actions (Jellema and Perrett, 2006; Perrett et al., 1989), the IPL infers the abstract goal or meaning of the observed actions (Fogassi et al., 2005; Hamilton, 2006), and the IFG and premotor area mediate the motor representation of the kinematic features of observed actions, and engagement in the action planning of mimicking behaviors (Hamilton, 2008). The IFG and pSTS are directly connected by the long segment of the arcuate fasciculus and indirectly connected by the anterior and posterior segments of the arcuate fasciculus via the IPL (Catani et al., 2005; Glasser and Rilling, 2008). Hamilton further proposed that the direct route between the IFG and pSTS allows the formation of direct associations from visual kinematic features in the pSTS to motor kinematic representations in the IFG, supporting mimicry behavior. The indirect route involves a two-stage procedure of goal emulation in the IPL and action planning in the IFG (Hamilton, 2008; Kilner et al., 2007). Spontaneous resting-state functional connectivity in the MNS has been documented in children aged 3 to 5 (Dai et al., 2019). A recent effective connectivity study showed that cooperative social interactions enhanced bidirectional effective connectivity between the pSTS and ventral premotor cortex, and effective connectivity from the pSTS and ventral premotor cortex to the superior parietal cortex (Arioli et al., 2018). Another effective connectivity study reported that volitional imitation of facial expressions enhanced bidirectional effective connectivity between the pSTS and IFG and between the IFG and IPL, and enhanced effective connectivity from the pSTS to the IPL (Sadeghi et al., 2022). A recent meta-analyses of fMRI studies using dynamic facial expressions indicated the involvement of only the right IFG (BA 9) and bilateral pSTS (Zinchenko et al., 2018), and another recent meta-analysis of facial expression processing, including studies using both static and dynamic facial expressions indicated the involvement of the ventral pathway, but not the IPL (Liu et al., 2021), which was consistent with the finding that macaques facial communicative visual inputs reach mirror neurons in area F5 via pathways that do not involve the parietal cortex (Ferrari et al., 2017).

The mentalizing network includes the IFG, anterior temporal lobe (aTL), temporoparietal junction (TPJ), dorsal and ventral medial prefrontal cortex (mPFC)/anterior cingulate cortex (ACC), and precuneus/posterior cingulate cortex (PCC) (Arioli et al., 2021; Redcay and Schilbach, 2019; Schurz et al., 2020). This network is involved in mentalizing or perspective-taking about the emotion of others, as part of the cognitive empathy process (Gonzalez-Liencres et al., 2013). Furthermore, the salience network, including the limbic regions (Seeley et al., 2007), together with the IFG, is involved in affective empathy (Walter, 2012). Therefore, individuals form representations of another’s feelings by sharing these feelings through embodied simulation (Gallese and Sinigaglia, 2011; Gonzalez-Liencres et al., 2013). These networks and cognitive affective processes are fundamental to emotional contagion and prosocial behaviors such as motor mimicry (Prochazkova and Kret, 2017).

However, previous neuroimaging studies have presented pre-recorded video stimuli to investigate the processing of dynamic facial expressions, with no potential for participants to interact with the observed agents making the expressions. Recent social neuroscientific studies have emphasized the importance of ecological validity in a face-to-face, real-time dyadic interactive design (Redcay and Schilbach, 2019; Schilbach et al., 2013; Shamay-Tsoory and Mendelsohn, 2019). In a previous study, we used a video camera relay system to allow participants to view live model performance or pre-recorded dynamic facial expressions. Our facial electromyogram (EMG) and subjective rating results showed that live facial expressions evoked stronger spontaneous facial mimicry and emotional contagion (Hsu et al., 2020). However, empirical evidence regarding neurocognitive processing of live dynamic facial expressions is lacking.

In the present study, we tested the effects of face-to-face live interaction via dynamic facial expressions on subjective emotion and facial mimicry in a population of 44 young female participants and two female models. We adapted the video relay system used in our previous study for the MR scanner environment (Fig. 1A). We recruited only female participants and models to control for sex effects in facial mimicry responses (Seibt et al., 2015). In the scanner, each participant viewed pre-recorded or live videos of positive (smiling) or negative (frowning) dynamic facial expressions, while multiecho echo-planar imaging (EPI) (Kundu et al., 2017) and EMG activity of the zygomaticus major (ZM) and corrugator supercilii (CS) were recorded. These procedures were followed by a functional localizer task and a rating task (Fig. 1B) to assess valence and arousal (Russell, 1980; Russell et al., 1989; Russell and Mehrabian, 1977).

**Figure 1.**
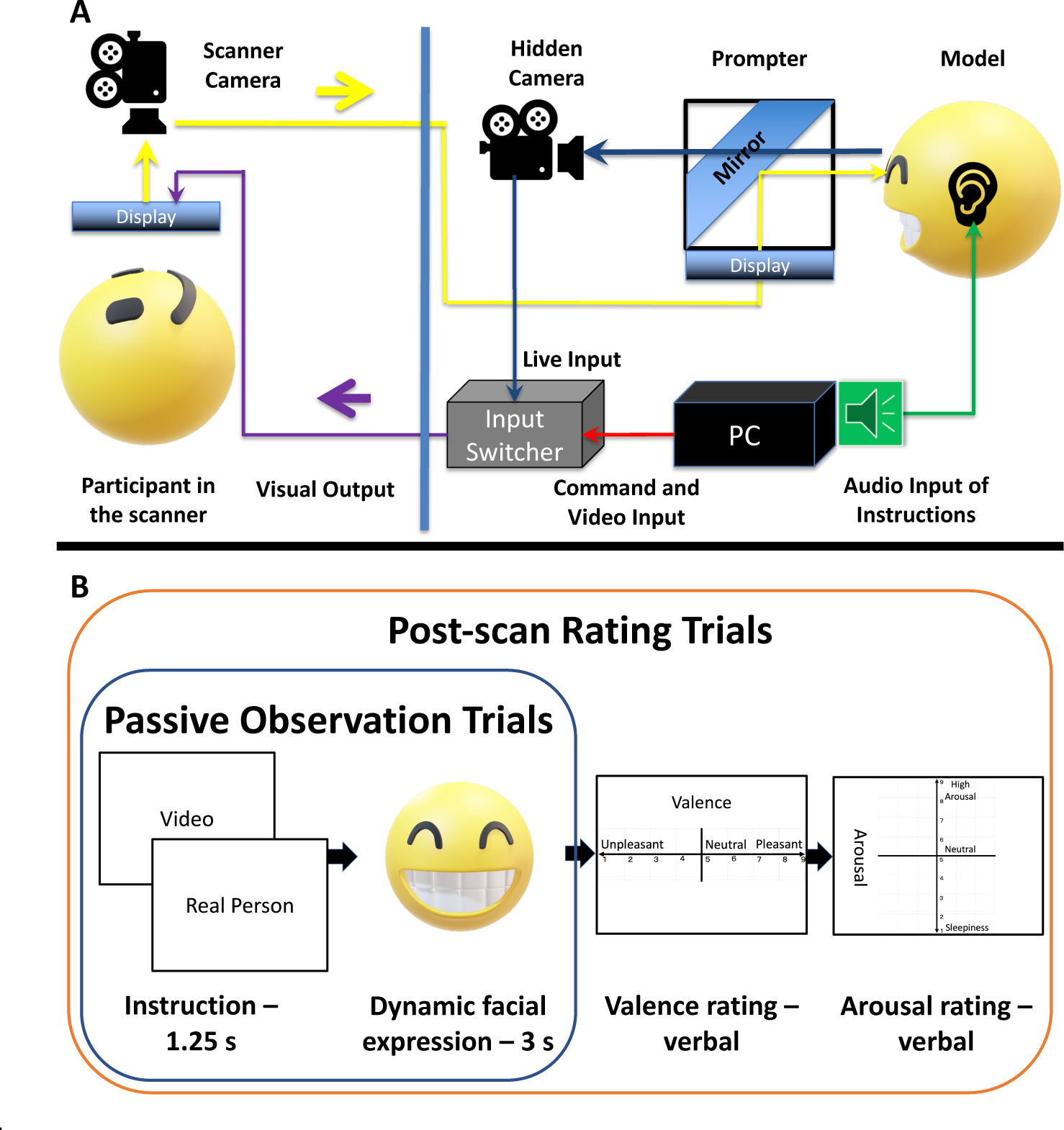
Experimental facilities (A) and paradigm (B). (A) The model outside the scanner faced a prompter. The prompter consisted of a horizontally positioned display and an obliquely positioned transparent mirror. The image of the model was captured by a camera hidden behind the mirror and sent to the input switcher (blue route) as a visual output option. The personal computer (PC) running the paradigm sent commands (red route) through a serial port to the input switcher, to determine the visual output for the participant to view (purple route). During video trials, the PC sent the instructions and pre-recorded video stimulus (red) to the input switcher, and the switcher sent only the input from the PC to the participant’s display (purple). During live trials, the switcher first relayed the instructions from the PC to the participant (red), and then, the PC transmitted the audio signal to the model via a pair of computer loudspeakers, instructing her to produce dynamic facial expressions (green route) while sending commands (red) to the input switcher to switch to the model’s live image input (blue) and to switch back after the duration of the stimulus (3 s). The participant in the scanner viewed the visual information on the display at the end of the scanner bore via a reflection in the mirror mounted on the head coil, while the participant’s image was video-recorded and live-relayed by a camera at the end of the scanner bore. The participant’s image was then sent to the prompter of the model (yellow route). (B) During passive observation trials, the participant was shown an instruction for either “Video” or “Real Person” followed by 3 s of pre-recorded or live dynamic facial expression, with the fixation cross lasting for the duration of the jittered interstimulus interval. In the post-scan rating trials, the participants was shown the instruction followed by the stimulus as in the passive observation trials, and were asked to report her valence and arousal ratings to the experimenter via an intercom inside the scanner room.

We expected to reproduce previous enhanced spontaneous mimicry and emotional contagion findings (Hsu et al., 2020) under live conditions in our EMG and rating data among female participants. Based on these findings, we further hypothesized that live interaction enhances mirroring and mentalizing, stimulating the MNS and mentalizing network into a more socially engaged state. Such a change in the brain activation state could be detected as a change in (1) the fMRI activity amplitude using univariate analysis, (2) a changed in the functional organization of neuronal units in a brain substrate or across the brain, presented as multivariate activation pattern change (Brooks et al., 2020; Weaverdyck et al., 2020), or (3) a change in connectivity within the brain network (Friston, 2011). We expected to detect enhanced activation using univariate analysis, or as a differential multivoxel activation pattern between live and pre-recorded conditions, as well as modulated functional connectivity in the MNS and mentalizing networks. We first examined the univariate activation patterns and defined functional region-of-interest (ROI) masks for effective connectivity analysis based on the univariate results. For substrates in the MNS and mentalizing networks that did not show univariate state changes, we further examined the possibility of multivariate activation pattern differences. The results of this study provide information on how the dynamics of the neural networks involved in social cognition differ between a “third-person” and a “second-person” approach (Redcay and Schilbach, 2019; Schilbach et al., 2013).

## 2. Methods

### 2.1 Participants

Fifty-one women from the Japanese cities of Kyoto and Osaka were recruited to this study in February and March 2021 via email lists, online advertisements, and flyers. This study was approved by the ethics committee of the Unit for Advanced Studies of the Human Mind, Kyoto University (approval number: 30-P-7), and performed in accordance with the ethical standards of the committee. Written informed consent was obtained from all participants prior to their participation in the study. All participants consented to the publication of their data, and to being recorded on video. All participants were rewarded monetarily. After excluding five participants who fell asleep in more than 3 of 5 runs of passive-viewing fMRI tasks (see section 2.6) and two who did not finish the fMRI tasks, 44 participants were included in the ratings and fMRI analysis, aged 22.82 ± 2.59 years [mean ± standard deviation (SD); range: 20-29 years]. EMG recordings failed for one of the 44 participants failed due to equipment setup error, whereby neither the scanner pulse nor experimental condition timing markers were stamped in the data and could not be reconstructed precisely, leaving 43 participants for the EMG and fMRI parametric modulation analyses.

To simulate the power of the interaction effect between emotion and presentation condition with extended participant numbers up to 55, we used the simr package (Green and MacLeod, 2016), based on zygomaticus major EMG linear mixed-effect model analysis from a previous study that used the same paradigm (Hsu et al., 2020). The simulation demonstrated that more than 17 participants were needed for the interaction effect of zygomaticus major EMG data to reach a power of 80%. No *a prior* power estimation was performed to determine the sample size required for the fMRI statistics of the passive viewing task, as the effect size of the presentation condition in this paradigm could not be approximated in previous neuroimaging studies. However, we opted for a sample size of 50, and finally included 44, which was in the top 20% of sample sizes in recent experimental fMRI studies (Szucs and Ioannidis, 2020). Using the NeuroPowerTools (Durnez et al., 2016), we used the statistical t-map of the functional localizer task adopted in a previous study (Sato et al., 2019). Power calculation showed that a sample size of 30 had a power of 0.85. NeuroPowerTools considers only peak inference, and the power for cluster inference was estimated under the assumption that there was no peak-level correction.

### 2.2 Materials

#### 2.2.1 Pre-recorded videos

As in a previous study (Hsu et al., 2020), two female models each recorded more than 40 video clips of positive (smiling) and negative (frowning) dynamic facial expressions. The appearance of the models was kept consistent between the video clips and live stimuli. Each clip lasted 3 s, and consisted of a neutral expression, gradual dynamic changes, and sustained maximal emotional facial expression, for 1 s each. The videos were saved and presented at a rate of 30 frames per second (fps). We used 50 smiling and 50 frowning clips for the passive viewing component; for each condition, two clips were used for the practice ratings, and four clips were used for the actual ratings. We randomly selected 10 clips per condition for each run of passive viewing.

#### 2.2.2 Facilities

The model outside the scanner faced a prompter with a concealed VIXIA HF R800 camera (Canon Inc., Tokyo, Japan). A camera in the scanner captured “live-relay images” of the participant to the model. The Presentation v21 software (Neurobehavioral Systems, Inc., Berkeley, CA, USA) was used to run the experiments on a Precision T3500 computer (Dell Inc., Round Rock, TX, USA) with the Windows 7 Professional operating system (Microsoft Corp., Redmond, WA, USA). Serial signals were sent to an SL-41C switcher (Imagenics Co., Ltd., Tokyo, Japan), to switch the visual input of the scanner screen between the computer and the model’s camera. Presentation software transmitted sounds to the models via a pair of computer loudspeakers, instructing them to produce dynamic facial expressions. The resolution (1280 × 960) and height-to-width ratio of the video output of the Presentation software and live-relay images were kept consistent by the switcher. The BrainAmp ExG MR amplifier (Brain Products, Munich, Germany) and BrainVision Recorder software (Brain Products) were used to record EMG from the ZM and CS of the participants.

### 2.3 Paradigm and procedures

As in a previous study (Hsu et al., 2020), we employed a 2 × 2 study design, with presentation conditions (video vs. live) and emotion conditions (positive vs. negative) as factors, resulting in four conditions. After providing written informed consent and the attachment with EMG electrodes, the participant was placed inside the scanner with a microphone. The participant and model engaged in audiovisual conversation for 3 min through the prompter and interphone. Subsequently, the structural T1 image was acquired.

#### 2.3.1 Passive Viewing Task

The participants fixated on a cross (mean inter-trial interval = 4475 ms; range: 2950 - 10150 ms) until they were shown instructions for either “Video” or “Real Person” for 1.25 s. During video trials, a pre-recorded video clip was presented immediately after the instructions. During live trials, the models performed dynamic facial expressions according to instructions provided across the loudspeaker. The screen displayed the fixation cross again after the facial expressions had been shown (Fig. 1B). The participants performed eight practice trials without data acquisition. Five functional task runs were performed, during which multi-echo EPI and EMG data were acquired. Each run included 10 passive viewing trials per condition and lasted 361.6 s (113 volumes). Due to input error in text files indicating the condition sequences, some runs comprised 11 “positive-live” trials and 9 “negative-live” trials. This phenomenon occurred in 2 of 5 runs for 14 participants, and 1 of 5 runs for another 14 participants. Between runs, data acquisition was stopped, and participants took breaks. A total of 200 trials were acquired per participant. For each participant, the sequence of conditions during the trial was pseudorandomized, and the presentation sequence of the pre-recorded videos per condition was randomized. The pseudorandomized presentation sequences and jittered inter-trial intervals were optimized using the optseq2 tool based on the cost function of the average variance reduction factor after 300,000 simulations (Dale, 1999). Both the models and participants were videotaped during data acquisition.

#### 2.3.2 Functional localizer task

The functional localizer task was modified from a previous study (Sato et al., 2019) using identical video clips of dynamic facial expressions (happy and angry) and dynamic mosaic images. Each clip lasted 1.52 s. In each epoch, the participant viewed eight clips of the same condition with the fixation cross presented for 1.68 s between clips. The onset of each clip was synchronized with the scanner pulse signal. Each epoch lasted for 25.6 s, and four epochs per condition were presented. The first fixation cross lasted for 35.2 s (11 TRs) until the onset of the first epoch. The inter-epoch interval and duration of the final fixation cross were both 19.2 s (6 TRs). This task lasted for 752 s (235 volumes).

#### 2.3.3 Subjective experiential valence and arousal ratings

After the functional localizer task, participants received instructions about and explanations for subjective experiential ratings with the Russell “affect grid” (Russell et al., 1989) shown on the screen: “When viewing the images, please rate the emotion that you feel. Please rate the emotion in two dimensions: (1) Pleasure-displeasure, which is about the quality of the emotion, and (2) arousal, which is about the energy of the emotion. Please speak out loud the number you find most suitable (1-9) in the sequence of ‘pleasure-displeasure’ and ‘arousal’.” Further explanation of valence and arousal using the affect grid followed: “The center of the square represents neutral emotion, which is neither positive nor negative; it is the ordinary emotion you experience. The right upper corner represents the emotion of excitement and happiness. The left lower corner represents the emotion of depression, sadness, or worry. The left upper corner represents stress and nervousness. The right lower corner represents tranquility and relaxation. We will first do four practice trials.” Subjective rating trials (Russell et al., 1989) began with the described passive-viewing component, and valence (pleasure-displeasure) and sequential arousal ratings were given verbally from inside the scanner after the subject had viewed dynamic facial expressions (Fig 1B). Four practice rating trials were followed by 16 test-rating trials. No fMRI or EMG data were acquired during this period. Subsequently, the participant was removed from the scanner and the electrodes were removed. Finally, the participant was monetarily rewarded and dismissed.

### 2.4 MRI data acquisition

Data were acquired using a 3T Siemens MAGNETOM Verio scanner with a 12-channel phased array coil. We acquired an MPRAGE scan with T_1_ weighted contrast [176 ascending sagittal slices with anterior-posterior (A/P) phase encoding direction; voxel size = 1 mm isotropic; field of view (FOV) = 256 mm×256 mm; repetition time (TR) = 2250 ms; echo time (TE) = 3.06 ms; acquisition time (TA) = 314 s; flip angle = 9°; GRAPPA in-plane acceleration factor = 2; complete brain coverage for the cerebrum, cerebellum, and brainstem]. After T_1_, we acquired five functional runs of T_2_* weighted echo planar sequence images of three echoes [40 interleaved axial slices with A/P phase encoding direction; voxel size = 3×3×3 mm; FOV = 192 mm×192 mm; TR = 3200 ms; TEs = 20, 37.29, and 54.58 ms; 113 volumes; TA = 361.6 s; flip angle = 90°; GRAPPA in-plane acceleration factor = 2; complete brain coverage for the cerebrum, cerebellum, and brainstem]. The MR sequence was compatible with the specific absorption rate requirement of BrainAmp ExG for simultaneous physiological-fMRI measurements (one pulse per slice, flip angle ≤ 90°, TR > 2000 ms, and ≤ 25 slices per 2000 ms). The same multi-echo sequence was used to acquire 235 volumes (TA = 752 s) for the functional localizer task.

### 2.5 Validation of live stimuli

We recorded and visually inspected the live performances of the models during the passive viewing trials to ensure valid performance of dynamic facial expressions. Three trials were excluded due to incorrect facial expression performance.

To determine whether live model performances could be more dynamic than those in the pre-recorded videoclips, we used OpenFace (Baltrušaitis et al., 2018, 2015) to quantify framewise amplitudes of facial action unit 4 (AU 4, brow lowerer, corresponding to the CS) and AU12 (lip corner puller, corresponding to the ZM) in video recordings of live model performances and pre-recorded clips (Ekman et al., 2002). All videos were recorded at 30 fps. The live model performances for one participant were not recorded, leaving data for 43 participants in this analysis. We calculated maximal dynamic change – the difference between the maximal amplitude in the third second (the 61^st^ frame to the final frame) and the minimal amplitude in the first second (the 1^st^ frame to the 30^th^ frame). We calculated amplitude differences of AU12 for positive trials, and those of AU4 for negative trials. Then, we calculated the mean differences between the trial-wise amplitude differences for positive-live, positive-video, negative-live, and negative-video conditions for each participant. Two group-level paired t-tests were performed to compare the participant-wise mean AU12 dynamic changes between positive-live and positive-video conditions and the participant-wise mean AU4 dynamic changes between negative-live and negative-video conditions. In positive conditions, the dynamic change for AU12 was significantly weaker for live performances (mean ± SD = 2.615 ± 0.489) than for pre-recorded clips (mean ± SD = 2.985 ± 0.460; difference = −0.370, standard error [SE] = 0.021, 95% confidence interval [CI] of difference = [−0.327, −0.413], *t_(42)_* = −17.504, *p* < 0.0001). In negative conditions, the dynamic change for AU4 was significantly weaker for live performances (mean ± SD = 1.467 ± 0.291) than for pre-recorded clips (mean ± SD = 2.024 ± 0.232; difference = −0.556, SE = 0.045, 95% CI of difference = [−0.464, −0.648], *t_(42)_* = −12.232, *p* < 0.0001).

### 2.6 Participant scanner performance check

We recorded and visually inspected video recordings of participants inside the scanner. Runs in which a participant closed her eyes for more than 50% of the time were excluded from EMG and fMRI data analyses. If three or more runs were excluded, the participant was completely excluded from EMG and fMRI data analysis. Five participants were excluded for this reason. Among the remaining 44 participants, four runs for eight participants and three runs for seven participants were included in the analyses. During the passive viewing task, 13 participants were fully awake (asleep for 0% of the run) in all 5 runs; four were fully awake in 4 of 5 runs; two were fully awake in 3 of 5 runs, 12 were fully awake in 2 of 5 runs, and eight were fully awake in 1 of 5 runs.

### 2.7 EMG data pre-processing

Raw EMG data were pre-processed using the BrainVision Analyzer 2 software. MR gradient artifacts were removed based on the pulse trigger, and data were down-sampled to 500 Hz. Data were then imported and pre-processed using the EEGLAB MATLAB toolbox v2019.1 (Swartz Center for Computational Neuroscience, San Diego, CA, USA). A high-pass filter was applied at 20 Hz. For each trial, the signal was detrended and the baseline was corrected for low frequency drift. The baseline for this step was defined as the mean of values from 3 s before to 1 s after stimulus onset, which was 2 s before and after the onset of the trial/instruction. For each signal, the absolute or rectified value plus one was natural log-transformed to correct for right-skewness of the raw data distribution.

### 2.8 Statistical analyses of behavioral data

Linear mixed-effects (LME) models, model comparisons, and influence diagnostics were performed using R software v4.2.1 (R Core Team. Vienna, Austria) with the *lme4* v1.1.30, *lmerTest* v3.1.3, *HLMdiag* v0.5.0, *emmeans* v1.8.0, *simr* v1.0.6 packages, and the optimizer BOBYQA. The dependent variables included valence rating, arousal rating, and the differences in EMG (ZM and CS) between the neutral (0-1 s after stimulus onset) and maximal phases (2.5-3.5 s after stimulus onset) of the dynamic facial expression. For each LME model, emotion condition, presentation condition, and their interaction were treated as fixed effects, with participant as a random factor. The reference level for the emotion condition was “Negative”, and that for the presentation condition was “Video”. Model complexity was increased stepwise in terms of the number of random effects; each model was compared with the previous (i.e., less complicated) model using an F-test [lmerTest::anova()] until the more complex model was no longer significantly superior to the simpler model. When the model estimation resulted in a singular fit, the model iteration process was considered complete, and the simpler model was selected as the final model. Initially, random intercepts for participants were included; the inclusion of by-subject random slopes for the effects of emotion condition, presentation condition, and their interaction term depended on the results of the model comparison process.

Homogeneity of residual variance is an *a priori* assumption to ensure valid inferences from LME models (Baayen and Milin, 2010; Loy and Hofmann, 2013). For LME model diagnostics, upward residual and influence analyses were performed using the HLMdiag package (Loy and Hofmann, 2014). Trials with an absolute standardized residual > 3, or Cook’s distance > 1.5 × the interquartile range (IQR) above the third quartile (*Q_3_* + 1.5 × *IQR*) were excluded. Then Cook’s distance was checked to exclude highly influential participants.

Satterthwaite’s formula was used to evaluate the degrees of freedom (df). The effect size of the semi-partial R^2^ (variance explained) of fixed effects was estimated using *r2glmm* 0.1.2 with the Kenward-Roger approach (Edwards et al., 2008; Jaeger et al., 2017; Nakagawa and Schielzeth, 2013). Effect sizes for semi-partial R^2^ values of regression models are considered small, medium, and large at 0.01, 0.09, and 0.25, respectively (Gignac and Szodorai, 2016). Interactions between emotion categories and presentation conditions (live vs. video) were evaluated by simple effects analysis using the *emmeans* function. Statistical significance was determined according to *p-*values for two-tailed tests.

### 2.9 fMRI data pre-processing

DICOM files were converted into 4D Nifti files using dcm2niix_afni v1.0.20181125. Multi-echo EPI pre-processing was performed using the tedana v0.0.8 library (https://doi.org/10.5281/zenodo.4725985) and SPM12 v7771 software (http://www.fil.ion.ucl.ac.uk/spm). Realignment parameters were estimated for the first echo volumes of all runs, with the participant’s first echo volume of the first functional run used as the reference. Slice timing was corrected toward the first slice (acquired at the mid-point of the TR) of each frame. Realignment parameters estimated using the first echo volumes were then applied to all echo volumes of the same run and resliced while retaining the original resolution (3 × 3 × 3 mm).

The TE-dependent tedana processing workflow (Kundu et al., 2013, 2012) (https://doi.org/10.5281/zenodo.5227302) was as follows. For each run, an initial mask was generated from the first echo using the compute_epi_mask function in the Nilearn package. An adaptive mask was then generated, in which each voxel value reflected the number of echoes with good data. A monoexponential model was fit to the data at each voxel using log-linear regression to estimate T2* and S0 maps. For each voxel, the value from the adaptive mask was used to determine which echoes would be used to estimate T2* and S0. Multi-echo data were then optimally combined using the t2s combination method (Posse et al., 1999). Global signal regression was applied to the multi-echo and optimally combined datasets. Principal component analysis followed by the Kundu component selection decision tree (Kundu et al., 2013) were applied to the optimally combined data for dimensionality reduction. Independent component analysis was performed to decompose the dimensionally reduced dataset. A series of TE-dependence metrics were calculated for each ICA component, including Kappa, Rho, and proportion of variance explained. Next, component selection was performed to identify blood oxygen level-dependent (BOLD, TE-dependent), non-BOLD (TE-independent), and uncertain (low-variance) components using the Kundu decision tree v2.5 (Kundu et al., 2013). T1c global signal regression was then applied to the data to remove spatially diffuse noise. This workflow used the NumPy (van der Walt et al., 2011), sciPy (http://www.scipy.org/), pandas (McKinney, 2010), scikit-learn (Pedregosa et al., 2011), Nilearn, NiBabel (http://doi.org/10.5281/zenodo.3233118), and the Dice similarity index libraries (Dice, 1945; Sørensen, 1948) in a conda environment with Python v3.6.12.

Using SPM12, we coregistered the structural image to the first frame of the denoised EPI of the first functional run, and segmented it into grey and white matter (Ashburner and Friston, 2005). Forward deformation parameters to the MNI152 space were estimated using geodesic shooting registration (Ashburner and Friston, 2011) to a geodesic shooting template provided in the Computational Anatomy Toolbox 12. Images were normalized using the 4^th^ degree B-Spline Interpolation algorithm in the original resolution (3 × 3 × 3 mm) and further smoothed with a Gaussian kernel of 6 mm full width at half maximum.

### 2.10 fMRI data analysis

#### 2.10.1 Univariate contrast

General linear model (GLM) and dynamic causal modelling (DCM) analyses were performed with SPM12. In the GLM analysis, the design matrix contained four psychological regressors of interest, “video-positive”, “video-negative”, “live-positive”, and “live-negative” conditions, specifying the onset of video/live dynamic facial expression. The duration of each trial was 3 s. Two additional psychological regressors specified the onsets of the instruction for “Video” or “Real Person”, lasting 1.25 s. We included six motion parameters as regressors of no interest. We applied a high-pass filter with a cutoff period of 128 s, and temporal autocorrelation was accounted for with the FAST option in SPM12 (Corbin et al., 2018). We performed analysis of variance (ANOVA) with two within-subject factors (presentation condition and emotion condition) and no between-subject factor. The partitioned error approach is recommended for SPM (Penny and Henson, 2006), in which first-level analyses produce contrast images, summarizing the effects for each subject. These images are entered as data into separate second-level models (“SPM Group Analysis,” 2019). Therefore, we calculated fixed effects across all runs for each subject. Contrasts of the average beta-images for all four conditions, as well as [positive > negative], [live > video] and [(live-positive > live-negative) > (video-positive > video-negative)] contrasts were calculated at the first level for each subject. At the group level, four random-effect, one-sample *t*-tests (*n* = 44) were performed to determine the average activation across four conditions, the main effects of emotion condition and presentation condition, as well as the interaction effect. We applied whole-brain cluster-level family-wise error (FWE)-corrected *p* < 0.05, using a cluster-defining threshold of *p* = 0.001 to control for multiple comparisons (Eklund et al., 2016; Flandin and Friston, 2017). The whole brain area was defined by a whole-brain inclusive mask defined by the Automated Anatomical Labeling (AAL) atlas (Tzourio-Mazoyer et al., 2002).

#### 2.10.2 Parametric modulation by facial mimicry

To investigate whether right IFG and pSTS activity modulated by the presentation conditions was associated with facial mimicry, we estimated the parametric effects of congruency-weighted contrasts of ZM and CS responses. Congruent EMG responses for positive expressions included contraction (positive response) of the ZM and relaxation (negative response) of the CS, and vice versa for negative expressions (McIntosh, 2006; McIntosh et al., 2006). For each trial, congruent ZM and CS responses were coded as 1 and incongruent responses were coded as 0. Trial-wise mimicking EMG responses (for positive expressions) were estimated as congruency-weighted contrasts of ZM and CS responses, and operationalized as [ZM congruency (1 or 0) ×run-wise normalized ZM responses] − [CS congruency (1 or 0) ×run-wise normalized CS responses]. In this GLM analysis, the design matrix contained one psychological regressor of interest: “dynamic facial expressions” including trials from all four conditions. The duration of each trial was 3 s. Trial-wise congruent EMG responses were used as the parametric modulator, including all trials in the runs included in fMRI analysis. Two additional psychological regressors specified the onset of the “Video” or “Real Person” instruction, which lasted 1.25 s. We included six motion parameters as regressors of no interest. We applied a high-pass filter with a cutoff period of 128 s, and the FAST option was used to account for the temporal autocorrelation (Corbin et al., 2018). Beta maps of the parametric effect across all runs for each subject were taken to the group level for the random-effect, one-sample *t*-tests (*n* = 43). We applied whole-brain cluster-level FWE-corrected *p* < 0.05, using a cluster-defining threshold of *p* = 0.001 to control for multiple comparisons. We specifically tested two MNS clusters significant for the [live > video] contrast, the right IFG and pSTS, using small volume correction (SVC) (Worsley et al., 1996) with functional masks for each cluster, applying peak-level FWE-corrected *p* < 0.05 to control for multiple comparisons.

#### 2.10.3 Laterality analysis

For laterality analysis, we compared the statistical t-maps of the GLM contrasts between the original and flipped images (Baciu et al., 2005; Kurth et al., 2015; Sato et al., 2019). During pre-processing, the realigned, slice-timing corrected, tedana TE-dependence processed, denoised functional images and the coregistered structural image were flipped along the x-axis. The flipped T1 image was then segmented (Ashburner and Friston, 2005) using a symmetric tissue probability map provided in the VBM8 toolbox. The forward deformation parameter estimation, normalization and smoothing procedures were identical to those of the original images (see Section 2.9). Using the pre-processed flipped functional images, we performed the same first-level GLM analyses for the presentation × emotion analysis, and the parametric modulatory effect of facial mimicry as described in Sections 2.10.1 and 2.10.2. Group-level random-effect paired *t*-tests were performed to compare contrast images of interest estimated by comparing the original images and flipped images, including the main effect of presentation condition, the main effect of emotion condition, the interaction effect between presentation and emotion condition, and the parametric modulatory effect of facial mimicry. An inclusive right-hemisphere mask was used to limit the group-level analysis to the right hemisphere. For each contrast, significant clusters more active in the original image than in the flipped image indicated right-dominant correlates, whereas the reverse indicated left-dominant correlates.

#### 2.10.4 Dynamic causal modeling

In the DCM analysis (Zeidman et al., 2019a, 2019b) (*n* = 44), we concatenated all runs of functional data per subject, and defined a single-run subject-level GLM model. The design matrix included one condition of interest, “faces”, including all trials of 3-s stimuli presentation, and one parametric regressor “presentation condition”, in which “live” and “video” trials were coded as 1 and −1. One additional psychological regressor of no interest specified the onset of the “Video” or “Real Person” instruction, which lasted 1.25 s. Other regressors of no interest included six motion parameters and run regressors to partial out the effects of different runs. The run regressors were created using the spm_fmri_concatenate.m function in SPM12. Two ROIs, the right pSTS and the IFG, were defined based on the group GLM result of the main effect of presentation condition [live > video] (Table 5). Another ROI, the right IPL, was defined based on the group GLM result of the average effect of all four facial stimulus conditions (Table S1). The time series eigenvariate of each ROI was extracted. In the full deterministic DCM model, we defined mutual bilinear connectivity as existent between three ROIs (A matrix). The “faces” condition was used as a psychological input to the pSTS (C matrix). The parametric modulator “presentation condition” (live vs. video) was set to modulate both between-region connections and self-connections (B matrix). The full DCM model of each subject was inverted for parameter estimation (Friston et al., 2003). The group means of each estimated full DCM model parameter were used as empirical priors to re-estimate the parameters, to avoid local maxima for higher accuracy (Friston et al., 2016). At the group level, a parametric empirical Bayes (PEB) model was set up to evaluate the posterior density of model parameters from single-subject model inversions. Bayesian model reduction (Friston et al., 2011) then provided the evidence and parameter distributions for all nested DCM models. Model evidence was compared among all nested models, and Bayesian model averaging (Trujillo-Barreto et al., 2004) yielded a weighted-average of the winning nested models. We report parameters, the posterior probability of which different from zero, is > 0.95.

#### 2.10.5 Functional localizer

GLM analysis was performed for the localizer task. We included 30 participants who were included in the task fMRI analysis and were awake during the localizer task (i.e., 14 of 44 participants included in the task fMRI analysis were asleep > 50% of the time during the localizer scan). Among the 30 participants, 17 were fully awake during the localizer task. The design matrix contained four psychological regressors of interest, “dynamic smiling expressions”, “dynamic frowning expressions”, “dynamic mosaic of smiling expressions”, and “dynamic mosaic of frowning expressions”, specifying the onset of the video clips. The duration of each trial was 1.52 s. We included six motion parameters as regressors of no interest, applied a high-pass filter with a cutoff period of 128 s to the first-level, fixed effect analysis, and accounted for the temporal autocorrelation using the FAST option in SPM12. The contrast [dynamic expressions > dynamic mosaics] was calculated at the first level for each subject. At the group level, we performed a random-effect, one-sample *t* test (*n* = 30) for the contrast. We applied whole-brain cluster-level FWE-corrected *p* < 0.05, using a cluster-defining threshold of *p* = 0.001.

#### 2.10.6 Multivariate pattern analysis

We used the Pattern Recognition for Neuroimaging Toolbox (PRoNTo) v2.1 (Schrouff et al., 2013b) for this analysis. We tested whether regions that are involved in dynamic facial expression processing, but showed no univariate amplitude difference between live vs. video conditions (i.e., bilateral mPFC/ACC and PCC, Fig. S3C), could be differentiated between the live vs. video condition based on voxel activation covariation patterns. The ROI was defined by a functional mask of the group-level GLM [dynamic face > dynamic mosaic] contrast in the functional localizer task, excluding the right IFG, right posterior temporal lobe and bilateral visual areas, which were also observed in the univariate result of the [live > video] contrast in the passive observation task. For all 44 participants, non-smoothed functional imaging data within the ROI mask were linearly detrended and mean-centered. A design matrix containing “video” and “live” conditions (3 s per trial) was used for volume selection. Under the assumption of a hemodynamic response delay of 6 s and dispersion of 6 s, volumes in which the BOLD signal corresponded to more than one condition were excluded from this analysis. A binary support vector machine (SVM) algorithm was implemented to classify volumes containing BOLD signals corresponding to “live” vs “video” conditions, and a 5-fold cross-validation of subjects (with 9, 9, 9, 9, and 8 subjects per fold) was employed to estimate the performance of the outer model loop. For each fold of the outer loop, the training set was further divided into training and testing sets using nested 5-fold cross-validation of blocks within each subject (the nested loop) to optimize the SVM soft-margin hyperparameter C (a trade-off between the decision boundary and misclassification in the SVM algorithm). The inner loop trained and tested the model with each hyperparameter value between 10^-3^ and 10^6^ in logarithmic steps. The parameter leading to the highest performance in the nested loop (balanced accuracy for classification) was then selected by the software in the outer loop (see PRoNTo 2.1 manual section 4.4.3). The result showed that 0.1 and 0.01 were the most frequently adopted C parameter. A total of 10,000 permutation were performed to create a null distribution by permuting the data labels, and to determine whether the balanced classification accuracy for the whole group was higher than that due to chance alone (mean accuracy of all permutations = 0.4999184). For the permutation analysis, SVM hyperparameter C optimization was limited to two values (0.1 and 0.01) to reduce computational time.

To infer the contributions of different anatomical regions to the differentiable multivariate patterns, we applied a whole-brain multiple kernel learning (MKL) algorithm using PRoNTo with the SimpleMKL package (Rakotomamonjy et al., 2008). MKL used *a priori* knowledge about brain anatomy, as defined by the AAL (Tzourio-Mazoyer et al., 2002), to segment the whole-brain multivariate pattern into regional patterns. The voxels of each *m* brain regions of the AAL constituted a basis kernel, and the whole-brain kernel was a linear combination of *m* basis kernels obtained by the contribution vector, *d_m_*, with a size of *m*, specifying the contribution of each basis kernel to the model. All elements of *d_m_* were ≥ 0, and the elements were summed to 1. SimpleMKL simplified the optimization problem to an SVM algorithm to solve the whole-brain kernel and gradient descents to obtain *d_m_*. Brain regions were then ranked according to *d_m_*, i.e., the percentage of the whole-brain normalized weight explained by each region (Schrouff et al., 2018, 2013a). We also opted for a 5-fold cross-validation of subjects to estimate model performance as the outer loop, and a nested 5-fold cross-validation of blocks within each subject as the nested loop for SVM hyperparameter C optimization. Hyperparameter C optimization was initially limited to a range of values between 10^-2^ and 10^5^ in logarithmic steps. A total of 1,000 permutations was performed to determine the significance and effect size of model performance. In these permutations, no nested loop for SVM hyperparameter C optimization was implemented to reduce computation time, and C was set to 0.1 because this value was selected by the software in all 5-fold cross-validations through hyperparameter optimization in the initial MKL analysis.

### 2.11 Reduced sample analysis

The inclusion of task fMRI runs of participants with eye closure time of up to 50% might be excessively liberal. Therefore, we performed additional analyses by including fMRI runs of only those participants with eye closure time < 25%. If ≥ 3 runs were excluded, the participant was completely excluded from data analysis. This approach reduced the quantity of data with non-task-associated-BOLD signals. However, it also significantly reduced the statistical power for univariate analysis, as well as the quantity of data available for the training and validation for MVPA. For the passive viewing task, data from 35 participants were analyzed. Of these, four runs for five participants and three runs for eleven participants were analyzed. The results are reported in the Supplementary Material Figs. S4 and S5 and Tables S5–9.

## 3. Results

### 3.1 Subjective ratings

#### 3.1.1 Valence ratings

The final LME model with the highest possible complexity for valence ratings included emotion condition, presentation condition, and their interaction as fixed effects, as well as random slopes for emotion condition, presentation condition and their interaction over “subject” as a random factor. After model diagnostics (Baayen and Milin, 2010), 638 data points for all 44 participants remained in the model. Significant main effects of emotion and presentation condition were detected. Participants felt more positive when viewing positive expressions than negative expressions, and when viewing the live expressions than the pre-recorded expressions. There was a significant interaction between emotion and presentation condition (Table 1, Fig. 2A). Simple effects analysis using the *emmeans* package showed that positive-live (mean ± SE = 7.05 ± 0.1282, df = 44.1, 95% CI = [6.79, 7.31]) evoked more positive emotion than negative-live conditions (mean ± SE = 3.39 ± 0.0968, df = 44.9, 95% CI = [3.20, 3.58]; difference = 3.66, SE = 0.198, df = 44.2, *t* = 18.501, *p* < 0.0001), and positive-video (mean ± SE = 6.09 ± 0.0873, df = 41.9, 95% CI = [5.91, 6.26]) evoked more positive emotion than negative-video conditions (mean ± SE = 3.71 ± 0.1152, df = 44.0, 95% CI = [3.48, 3.94]; difference = 2.38, SE = 0.169, df = 43.9, *t* = 14.061, *p* < 0.0001). Positive-live stimuli evoked stronger positive emotions than positive-video stimuli (estimate = 0.959, SE = 0.110, df = 43.8, *t* = 8.703, *p* < 0.0001), whereas negative-live stimuli evoked stronger negative emotions than negative-video stimuli (estimate = −0.317, SE = 0.085, df = 42.5, *t* = −3.733, *p* = 0.0006).

**Table 1.**
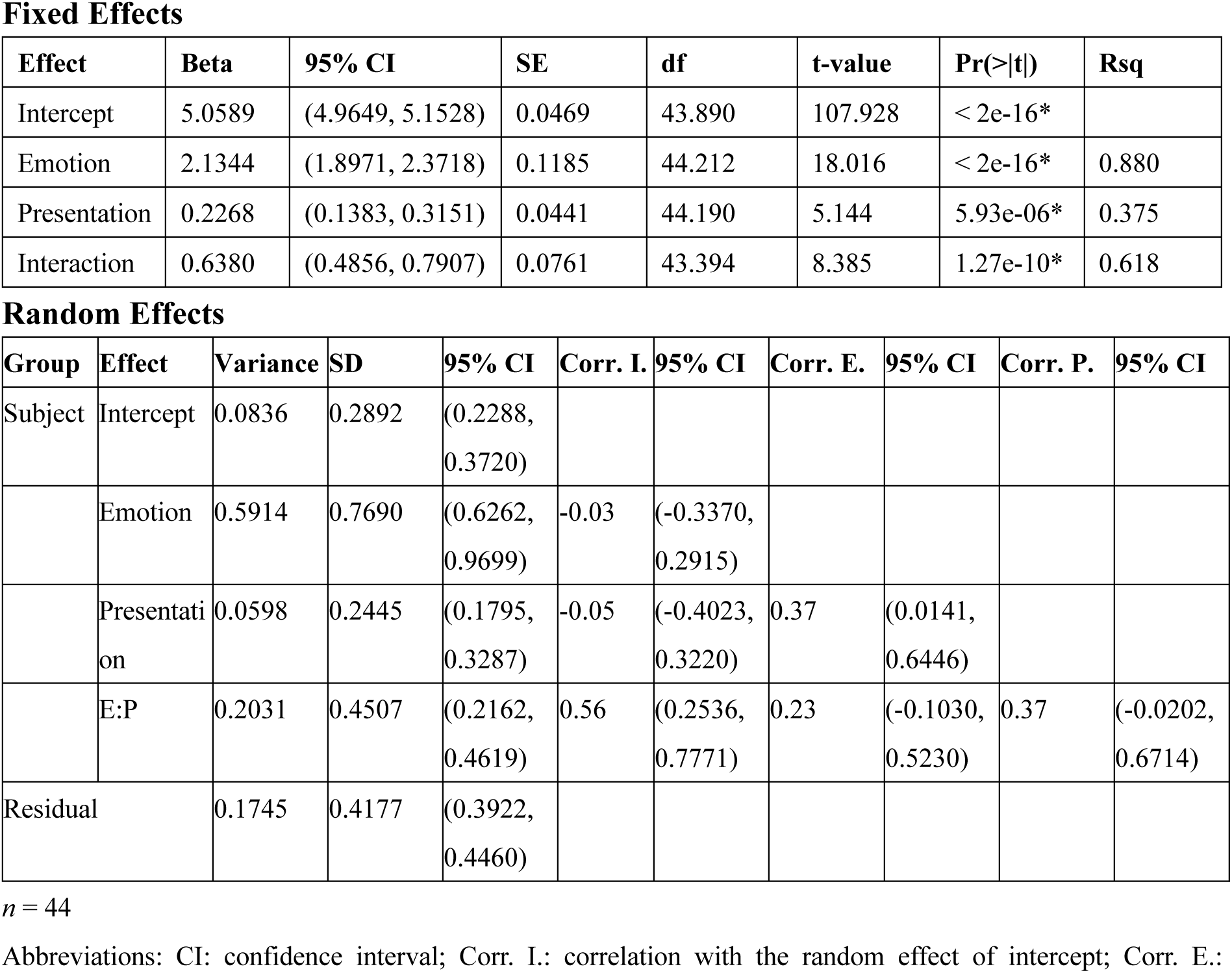
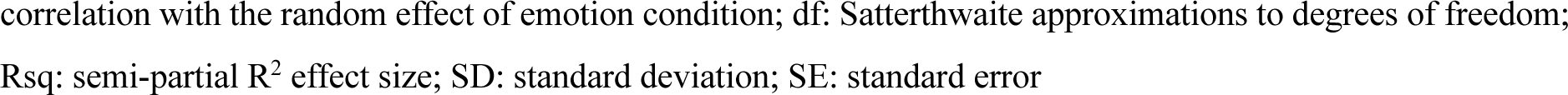
Statistical summary of valence ratings Fixed Effects

**Figure 2.**
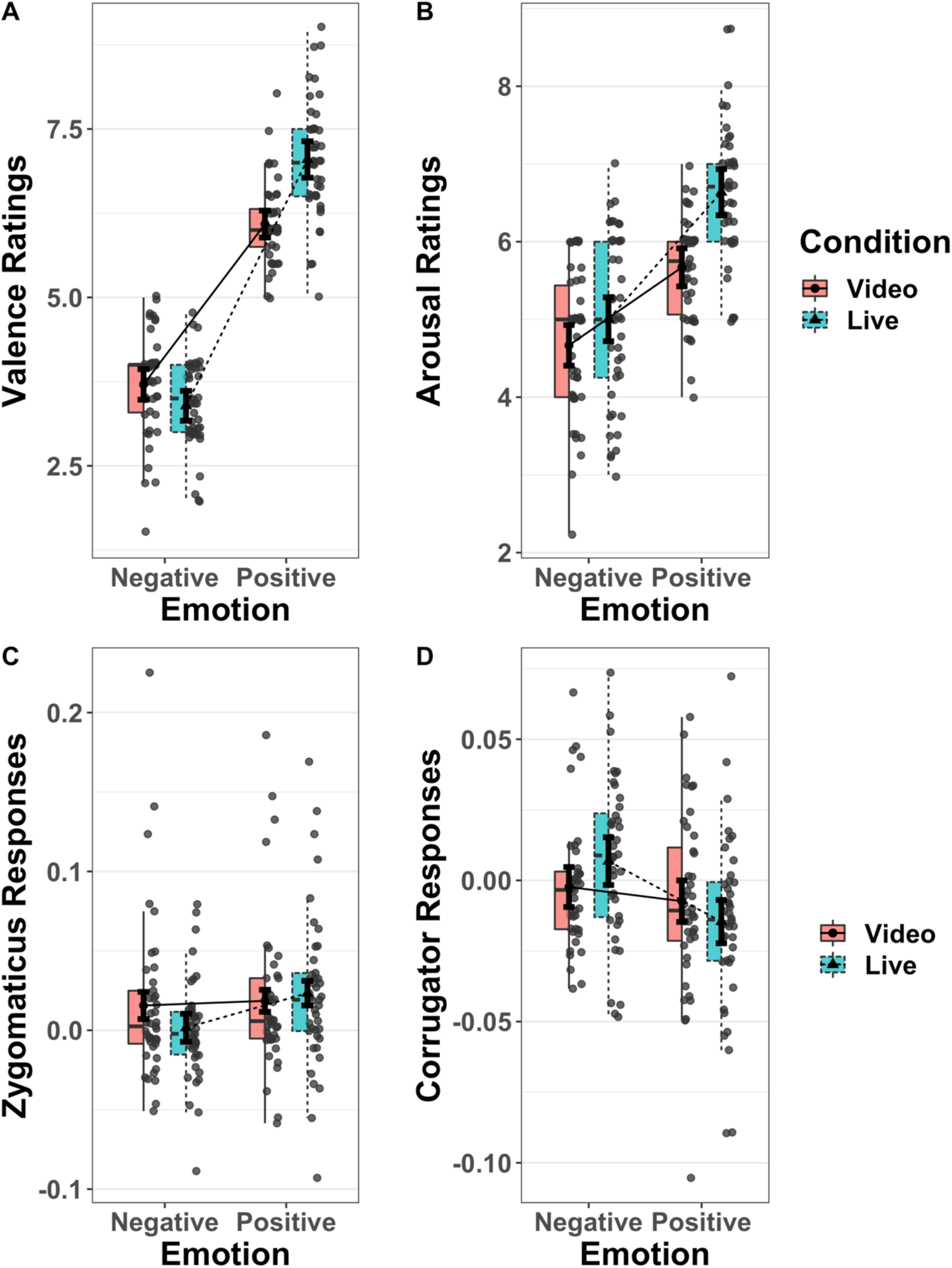
Interaction between presentation and emotion conditions on ratings and facial muscular reactions. (A) Valence ratings (*n* = 44). (B) Arousal ratings (*n* = 42). (C) Zygomaticus responses (*n* = 41). (D) Corrugator responses (*n* = 40). In all cases, significant interactions were observed between emotion conditions (positive vs. negative) and presentation conditions (video vs. live). For each condition, the right half shows the scattered dots of participant-wise mean values. Filled dot (video) and triangle (live) indicate group mean values, error bars indicate within-subject standard error. The box on the left half defines the median, along with the first and third quartiles of the distribution, with upper and lower whiskers extending from the hinge to the most extreme value, not exceeding than 1.5 × the interquartile range (IQR) from the hinge.

#### 3.1.2 Arousal ratings

The final LME model for arousal ratings included random slopes for emotion conditions and presentation conditions and their interaction over “subject” as a random factor. After model diagnostics, 602 data points of 42 participants remained in the model. There were significant main effects of the emotion condition and presentation condition, as well as their interaction. Positive conditions were rated as more arousing than negative conditions, and live conditions were rated as more arousing than video conditions (Table 2, Fig. 2B). Simple effects analysis showed that participants felt greater arousal in response to positive-live (mean ± SE = 6.64 ± 0.1372, df = 42.4, 95% CI = [6.36, 6.91]) than negative-live conditions (mean ± SE = 5.00 ± 0.1654, df = 41.9, 95% CI = [4.67, 5.34]; difference = 1.63, SE = 0.225, df = 42.3, *t =* 7.259, *p* < 0.0001) and greater arousal in response to positive-video (mean ± SE = 5.67 ± 0.0986, df = 41.4, 95% CI = [5.47, 5.87]) than negative-video conditions (mean ± SE = 4.67 ± 0.1410, df = 41.5, 95% CI = [4.38, 4.95]; difference = 1.00, SE = 0.191, df = 42.3, *t* = 5.252, *p* < 0.0001). Participants also felt greater arousal in response to positive-live than positive-video stimuli (difference = 0.967, SE = 0.1285, df = 41.7, *t* = 7.550, *p* < 0.0001), and more aroused to negative-live than negative-video stimuli (difference = 0.335, SE = 0.130, df = 41.2, *t* = 2.587, *p* = 0.0133).

**Table 2.** Statistical summary of arousal ratings Fixed Effects

| Effect | Beta | 95% CI | SE | df | t-value | Pr(>\|t\|) | Rsqr |
| --- | --- | --- | --- | --- | --- | --- | --- |
| Intercept | 5.4941 | (5.3432, 5.6454) | 0.0753 | 41.232 | 72.987 | < 2e-16* |  |
| Emotion | 0.9325 | (0.6615, 1.2032) | 0.1351 | 42.270 | 6.904 | 1.93e-08* | 0.530 |
| Presentation | 0.4605 | (0.3219, 0.5988) | 0.0689 | 39.972 | 6.682 | 5.24e-08* | 0.527 |
| Interaction | 0.3160 | (0.1473, 0.4856) | 0.0843 | 41.749 | 3.750 | 0.000538* | 0.252 |

Random Effects
| Group | Effect | Variance | SD | 95% CI | Corr. I. | 95% CI | Corr. E. | 95% CI | Corr. P. | 95% CI |
| --- | --- | --- | --- | --- | --- | --- | --- | --- | --- | --- |
| Subject | Intercept | 0.2164 | 0.4652 | (0.3706, 0.5974) |  |  |  |  |  |  |
|  | Emotion | 0.7228 | 0.8502 | (0.6866, 1.0814) | -0.35 | (-0.6000, -0.0313) |  |  |  |  |
|  | Presentation | 0.1566 | 0.3958 | (0.2989, 0.5257) | 0.39 | (0.0331, 0.6603) | 0.09 | (-0.2557, 0.4172) |  |  |
|  | E:P | 0.2127 | 0.4612 | (0.3378, 0.6224) | -0.07 | (-0.4287, 0.2992) | 0.28 | (-0.0886, 0.5805) | -0.01 | (-0.3957, 0.3762) |
| Residual |  | 0.2882 | 0.5369 | (0.5030, 0.5747) |  |  |  |  |  |  |

$n = 42$
Abbreviations: CI: confidence interval; Corr. I.: correlation with the random effect of intercept; Corr. E.: correlation with the random effect of emotion; df: Satterthwaite approximations to degrees of freedom; Rsq: semi-partial $R^2$ effect size; SD: standard deviation; SE: standard error

### 3.2 Facial EMG

#### 3.2.1 Zygomaticus major

The final LME model for the ZM responses included the random slope of emotion and presentation condition over the random factor, “subject”. During model diagnostics, 6574 data points for 41 participants remained in the model. The main effect of emotions was significant. ZM reactions were significantly stronger in the positive condition than in the negative condition. For the significant interactions between emotion and presentation conditions (Table 3, Fig. 2C), simple effects analysis showed that ZM reactions were stronger for positive-live (mean ± SE = 0.02336 ± 0.00719, df = 44.4, 95% CI = [0.00888, 0.0378]) than negative-live conditions (mean ± SE = 0.00158 ± 0.00561, df = 46.5, 95% CI = [−0.009718, 0.0129]; difference = 0.02179, SE = 0.00544, df = 77.8, *t* = 4.002, *p* = 0.0001), whereas there was no difference in ZM response between positive-video (mean ± SE = 0.01852 ± 0.00821, df = 42.8, 95% CI = [0.00197, 0.0351]) and negative-video conditions (mean ± SE = 0.01561 ± 0.00718, df = 43.2, 95% CI = [0.00112, 0.0301]; difference = 0.00292, SE = 0.00541, df = 75.8, *t* = 0.539, *p* = 0.5914). ZM contractions did not differ between positive-live and positive-video stimuli (difference = 0.00484, SE = 0.00477, df = 97.1, *t* = 1.014, *p* = .3130), whereas negative-live stimuli evoked more ZM relaxations than negative-video stimuli (difference = −0.01403, SE = 0.00480, df = 99.3, *t* = −2.924, *p* = 0.0043).

**Table 3.** Statistical summary of zygomaticus major reactions Fixed Effects

| Effect | Beta | 95% CI | SE | df | t-value | Pr(>\|t\|) | Rsqr |
| --- | --- | --- | --- | --- | --- | --- | --- |
| Intercept | 1.477e-02 | (0.0021, 0.0274) | 6.292e-03 | 40.65 | 2.347 | 0.02388* |  |
| Emotion | 8.734e-03 | (0.0023, 0.0153) | 3.224e-03 | 38.29 | 2.709 | 0.01003* | 0.161 |
| Presentation | -3.249e-03 | (-0.0087, 0.0021) | 2.668e-03 | 38.03 | -1.218 | 0.23069 | 0.038 |
| Interaction | 9.436e-03 | (0.0037, 0.0152) | 2.944e-03 | 6463 | 3.205 | 0.00136* | 0.002 |

**Random Effects**
| Group | Effect | Variance | SD | SD 95% CI | Corr. I. | Corr. 95% CI | Corr. E. | Corr. 95% CI |
| --- | --- | --- | --- | --- | --- | --- | --- | --- |
| Subject | Intercept | 0.0015318 | 0.0391 | (0.0315, 0.0489) |  |  |  |  |
|  | Emotion | 0.0002456 | 0.0157 | (0.0115, 0.0158) | 0.43 | (0.0219, 0.4276) |  |  |
|  | Presentation | 0.0001127 | 0.0106 | (0.0032, 0.0171) | -0.63 | (-0.7269, -0.1523) | 0.26 | (-0.4003, 0.3164) |
| Residual |  | 0.0142216 | 0.1193 | (0.1172, 0.1213) |  |  |  |  |

$n = 41$
Abbreviations: CI: confidence interval; Corr. I.: correlation with the random effect of intercept; Corr. E.: correlation with the random effect of emotion; df: Satterthwaite approximations to degrees of freedom; Rsqr: semi-partial $R^2$ effect size; SD: standard deviation; SE: standard error

#### 3.2.2 Corrugator supercilii

The final model for the CS responses included only the random slope for emotion conditions. After model diagnostics, 6477 data points for 40 participants remained in the model. With regard to the main effect of emotion conditions, the CS was more relaxed in the positive condition. There were significant interactions between emotion and presentation conditions (Table 4, Fig. 2D). Simple effects analysis showed stronger CS responses in the negative-live (mean ± SE = 0.00682 ± 0.00410, df = 47.3, 95% CI = [−0.00143, 0.01507]) than in the positive-live condition (mean ± SE = −0.01459 ± 0.00478, df = 45.3, 95% CI = [−0.02420, 0.00497]; difference = 0.02141, SE = 0.00522, df = 66.4, *t* = 4.104, *p* = 0.0001), whereas no difference was detected in CS responses between positive-video (mean ± SE = −0.00734 ± 0.00491, df = 45.0, 95% CI = [−0.01723, 0.00254]) and negative-video conditions (mean ± SE = −0.00234 ± 0.00364, df = 46.2, 95% CI = [−0.00966, 0.00499]; estimate = −0.00501, SE = 0.00519, df = 64.9, *t* = 0.9651, *p* = 0.3379). There was no difference in CS responses between positive-live and positive-video conditions (difference = −0.00724, SE = 0.00426, df = 87.7, *t* = −1.701, *p* = 0.0925), whereas negative-live stimuli evoked more CS contraction than negative-video stimuli (estimate = 0.00916, SE = 0.00436, df = 96.0, *t* = 2.101, *p* = 0.0382).

**Table 4.**
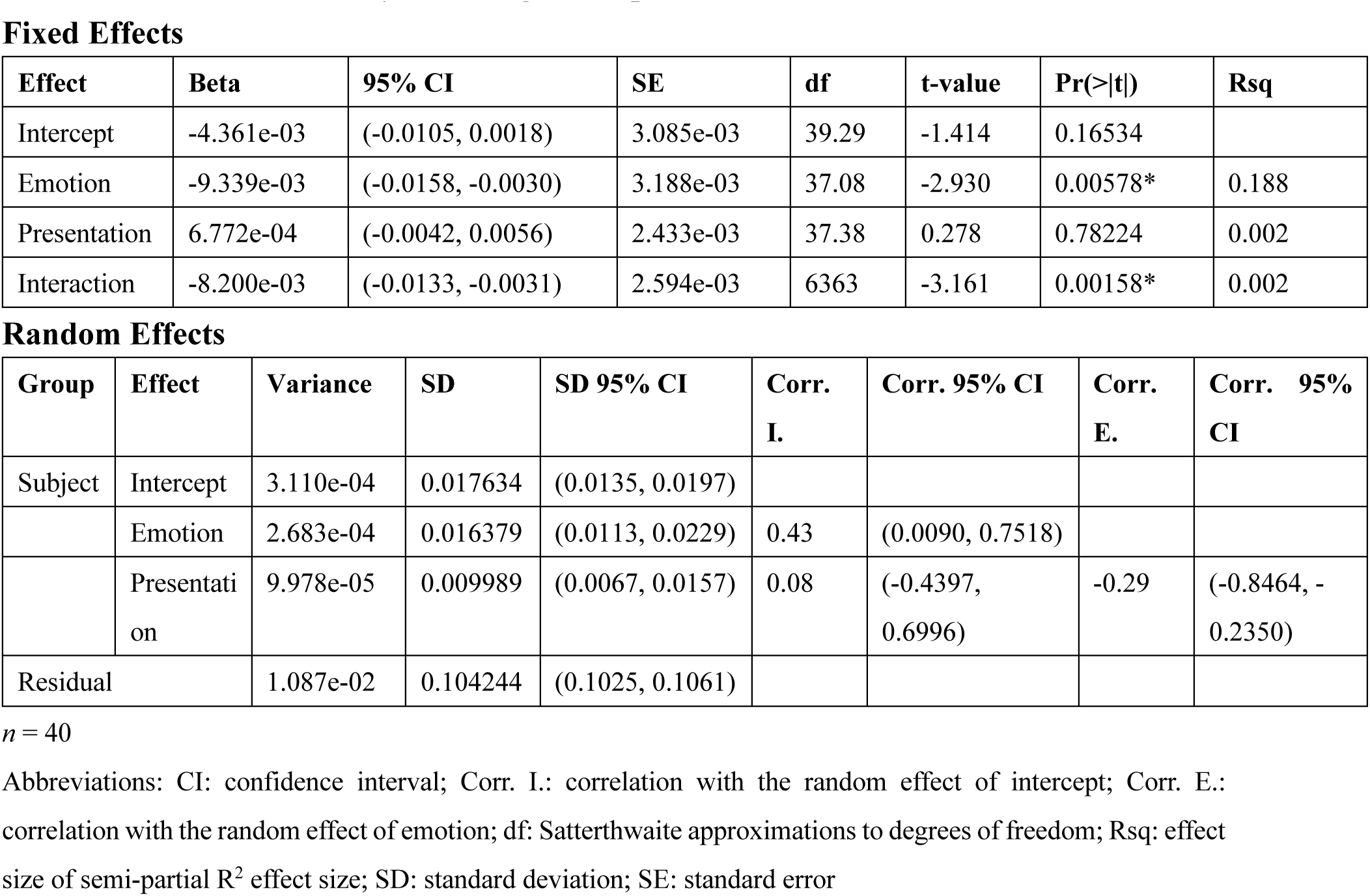
Statistical summary of corrugator supercilii reactions Fixed Effects

### 3.3 fMRI Results

#### 3.3.1 Univariate contrast

The univariate average activation pattern of the four conditions in the passive viewing task showed extensive bilateral IFG, supplementary motor area, precentral gyrus, supramarginal gyrus, IPL, hippocampus, and amygdala engagement (Supplementary Table S1, Fig. S1).

For the main effect of the presentation condition, the contrast [live > video] was associated with activities in the MNS – bilateral pSTS and right IFG – as well as right amygdala, right fusiform gyrus, and left visual cortex (Fig. 3A, B). The contrast [video > live] showed the right fusiform and lingual gyrus (Supplementary Fig. S2B). For the main effect of the emotion condition, the contrast [positive > negative] showed right IFG pars orbitalis and triangularis, and left occipital poles (Supplementary Fig. S2C), whereas the contrast [negative > positive] showed the right occipital pole (Supplementary Fig. S2D). There was no significant neural correlate for the interaction contrast of either direction (Table 5).

**Table 5.** fMRI GLM results

| H | Regions | Cluster size | p (FWE-corr)* | T | B.A. | [x, y, z] |
| --- | --- | --- | --- | --- | --- | --- |
| **Live > Video** |  |  |  |  |  |  |
| R | Posterior middle temporal gyrus | 1135 | <1e-40 | 10.50 | 39 | 54 -58 8 |
|  | Lingual gyrus |  |  | 9.25 | 18 | 18 -88 -7 |
|  | Posterior inferior temporal gyrus |  |  | 7.51 | 37 | 51 -67 -4 |
| L | Lingual gyrus | 142 | 4.0817e-05 | 7.32 | 17 | -15 -94 -7 |
| L | Posterior middle temporal gyrus | 384 | 1.7931e-10 | 6.88 | 39 | -57 -61 8 |
|  | Inferior occipital gyrus |  |  | 6.42 | 19 | -42 -82 -7 |
|  | Posterior superior temporal gyrus |  |  | 5.28 | 21 | -48 -52 8 |
| R | Middle frontal gyrus | 248 | 1.1157e-07 | 5.35 | 6 | 48 -1 53 |
|  | Precentral gyrus |  |  | 5.22 | 8 | 51 5 41 |
|  | IFG, pars triangularis |  |  | 4.67 | 45 | 45 23 20 |
| R | Amygdala | 59 | 0.0135 | 5.13 |  | 21 -7 -16 |
| R | Fusiform gyrus | 46 | 0.0407 | 5.06 | 20 | 39 -7 -31 |
|  | Fusiform gyrus |  |  | 5.02 | 36 | 27 -4 -37 |
| **Video > Live** |  |  |  |  |  |  |
| R | Lingual gyrus | 53 | 0.0223 | 5.90 | 18 | 15 -73 -4 |
| R | Fusiform gyrus | 114 | 2.4334e-04 | 5.83 | 37 | 30 -52 -10 |
|  | Fusiform gyrus |  |  | 5.55 | 37 | 27 -40 -19 |
|  | Parahippocampal gyrus |  |  | 4.90 | 37 | 36 -40 -7 |
| **Positive > Negative** |  |  |  |  |  |  |
| L | Occipital pole | 80 | 0.0041 | 5.37 | 17 | -24 -100 8 |
| R | IFG, pars orbitalis | 82 | 0.0036 | 4.15 | 47 | 51 11 5 |
|  | IFG, pars triangularis |  |  | 4.13 | 45 | 48 20 5 |
|  | IFG, pars triangularis |  |  | 4.08 | 45 | 60 20 2 |
| **Negative > Positive** |  |  |  |  |  |  |
| R | Occipital pole | 93 | 0.0017 | 5.18 | 18 | 27 -88 -13 |
| **Positive Correlation with Congruency-weighted Zygomaticus minus Corrugator Activities (Whole-brain)** |  |  |  |  |  |  |
| R | Inferior frontal gyrus | 167 | 6.6670e-05 | 4.93 | 9 | 42 -7 38 |
|  | Precentral gyrus |  |  | 4.50 | 4 | 57 -4 38 |
|  | Precentral gyrus |  |  | 4.22 | 4 | 42 -19 41 |
| R | Superior occipital gyrus | 77 | 0.0098 | 4.89 | 18 | 21 -91 8 |
| R | Posterior MTG | 68 | 0.0176 | 4.74 | 22 | 69 -43 2 |
|  | Posterior MTG |  |  | 4.13 | 21 | 54 -52 5 |
| L | Middle occipital gyrus | 85 | 0.0059 | 4.26 | 18 | -21 -97 5 |
|  | Middle occipital gyru |  |  | 3.85 | 18 | -15 -103 -1 |
| **Positive Correlation with Congruency-weighted Zygomaticus minus Corrugator Activities (Small Volume Correction)** |  |  |  |  |  |  |
| R | IFG | 4 | 0.0472 | 3.61 | 9 | 57 5 35 |
| R | Posterior MTG | 51 | 0.0253 | 4.45 | 22 | 69 -43 5 |
| * Cluster-level FWE-corrected for the whole-brain, cluster-forming threshold: uncorrected $p = .001$. Peak-level FWE-corrected for the small volume correction.<br>** Multiple peaks were listed for one cluster if the cluster spans across both hemispheres.<br>*** $n = 44$, except for the parametric modulation where $N = 43$.<br>Abbreviations: IFG = Inferior frontal gyrus; MTG = middle temporal gyrus; STS = superior temporal sulcus; H = hemisphere; L = left; R = right; $p$ = $p$-value; $T$ = $T$-value; B.A. = Brodmann area; $x, y, z$ = MNI coordinates |  |  |  |  |  |  |

**Figure 3.**
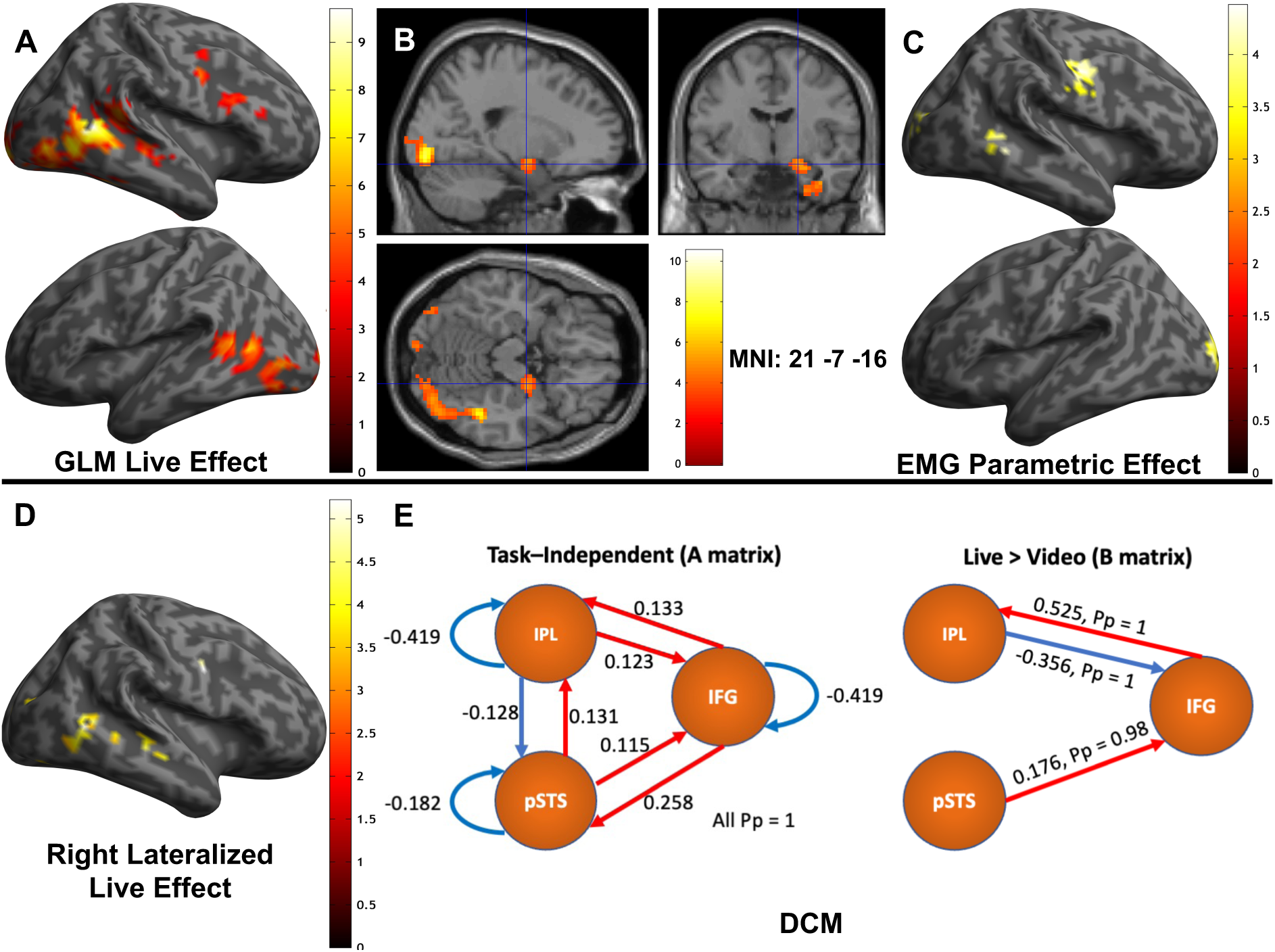
Univariate fMRI and dynamic causal modeling (DCM) results. (A) Statistical parametric maps of the main effects of presentation condition: [Live > Video]. Upper panel: Lateral view of the right hemisphere showing activations in the right inferior frontal gyrus (IFG) and posterior superior temporal sulcus (pSTS). Lower panel: Lateral view of the left hemisphere showing activation in the left pSTS (*n* = 44). (B) Section views with the crosshair (MNI = [21 −7 −16]), highlighting the right amygdala in the main effect of presentation condition [Live > Video]. (C) Statistical parametric maps of the parametric effect of congruency-weighted contrast of ZM and CS activities (*n* = 43). Upper panel: Lateral view of the right hemisphere showing activation in the right IFG, pre- and postcentral gyri and posterior superior and middle temporal gyri. Lower panel: Lateral view of the left hemisphere showing activation in the left occipital pole. (D) Statistical parametric maps of significant right dominant clusters for the main effect of presentation condition: [Live > Video], with right lateral view of clusters in the right mid- and pSTS, premotor area, precentral gyrus and visual cortex. (E) DCM results of right MNS – IFG, pSTS, and inferior parietal lobule (IPL). The circles indicate three regions of interest (ROIs). Red and blue lines indicate positive and negative effective connectivity, respectively. Numbers indicate connectivity strength (in Hz). Left panel: Task-independent baseline connectivity, estimated in the DCM as the A matrix. Right panel: Task modulatory effective connectivity, estimated in the DCM as the B matrix. In the design matrix, the parametric modulatory effect of live and video conditions were dummy-coded as 1 and −1, respectively (*n* = 44). Pp: posterior probability of parameter > 0.

#### 3.3.2 Parametric modulation by facial mimicry

At the whole-brain level, congruent EMG responses were correlated with activity in the right IFG, premotor cortex, precentral gyrus, posterior temporal lobe, and bilateral occipital poles. The small volume correction (Worsley et al., 1996) was significant for the right IFG and premotor cortex (with the peak located in IFG Brodmann area 9) and pSTS clusters, which also showed the main effect of presentation condition [live > video] (Table 5, Fig. 3C).

#### 3.3.3 Laterality analysis

For the main effect of presentation conditions, the contrast [live > video] showed right-dominant activation [right > left] in the mid- and pSTS, premotor area, precentral gyrus and visual cortex (Fig. 3D), whereas a different subregion in the visual cortex showed left-dominant activation (Table 6). There was no significant laterality for results of the main effect of emotion conditions, the interaction effect between two factors, or the parametric modulatory effect of facial mimicry.

#### 3.3.4 Dynamic causal modeling

Group DCM results (Fig. 3E) showed that baseline effective connectivity consisted of positive effective connectivity between all pairs of nodes except for the negative effective connectivity from the right IPL to the right pSTS. The modulatory effect of live over video conditions consisted of the positive effective connectivity from the right pSTS to the right IFG (0.176 Hz, Pp = 0.98), positive effective connectivity from the right IFG to the right IPL (0.525 Hz, Pp = 1), and negative effective connectivity from the right IPL to the right IFG (−0.356 Hz, Pp = 1).

#### 3.3.5 Functional localizer

The contrast [dynamic face > dynamic mosaic] was compatible with the results of a previous study using more stimulus blocks (Sato et al., 2019). Significant clusters included the bilateral amygdala, right posterior temporal cortex including the fusiform gyrus, and the mentalizing network including the aTL, mPFC, and PCC (Supplementary Table S2, Fig. S3). A functional mask containing the significant clusters of the bilateral mPFC and PCC was created for multivariate pattern analysis (MVPA, Fig. S3C).

#### 3.3.6 Multivariate pattern analysis

We further examined whether the presentation condition modulated regions involved in dynamic facial expression processing but found no amplitude difference between live vs. video conditions (i.e., bilateral mPFC and PCC, Fig. S3C) in forms of the multivariate voxel activation pattern. The binary SVM algorithm yielded a group-level classification accuracy of 54.94% (area under the receiver operating characteristic curve (AUC) = 0.56, positive predictive values for classes [live, video] = [56.58%, 53.96%]). A permutation test (Golland and Fischl, 2003), with 10,000 repetitions, showed that the accuracy was significantly higher than the 50% of the chance level (*p* = 9.999×10^-5^).

For the whole-brain MKL, the whole-brain group-level classification accuracy was 55.23% (AUC = 0.55, positive predictive values for classes [live, video] = [57.06%, 54.15%]). A permutation test with 1,000 repetitions yielded *p* = 9.99×10^-4^. The ranking of AAL ROIs that contributed > 1% to the whole-brain weight included the right ACC (50.79%) and right putamen (36.01%).

## 4. Discussion

### 4.1 Study summary and general univariate activation patterns

Using a video camera relay system, we investigated the effects of live interaction on subjective experiential, muscular, and neural networks that are important for social cognition. Under the live presentation condition, positive expressions were rated as more positive and arousing, and elicited stronger congruent motor responses in the ZM and CS (Tables 1-4, Fig. 2). These results are consistent with those of our previous behavioral study (Hsu et al., 2020), and indicate that live interactions enhance spontaneous facial mimicry and emotional contagion.

In the fMRI data, passive observations of both pre-recorded and live dynamic facial expressions involved brain regions classically associated with facial expression processing, such as the MNS, which includes the bilateral frontoparietal network and right posterior temporal lobe (Hamilton, 2008), the mentalizing network including the anterior cingulate cortex (BA 32), the motor system including the bilateral motor cortex and medial supplementary motor area (BA 6), and the salience network including the bilateral amygdala, hippocampus, right insula and right thalamus (Seeley et al., 2007) (Supplementary Table S1, Fig. S1). The whole-brain activation pattern was compatible with the findings of previous neuroimaging studies that had investigated facial expression processing (see the Introduction), and confirm participant engagement in the passive observation task.

### 4.2 Live effects modulated the MNS in univariate activation and connectivity

The main effect of presentation condition in the fMRI data showed stronger activation in the right MNS (pSTS, IFG and premotor cortex) when the participant observed live performances (Table 5, Fig. 3A, B). Activity in the right pSTS, IFG, premotor cortex and precentral gyrus was parametrically correlated with measures of spontaneous facial mimicry (congruent ZM and CS responses, Table 5, Fig. 3C). The live effect also modulated effective connectivity within the right MNS (enhanced from pSTS to IFG and from IFG to IPL, and reduced from IPL to IFG, Fig. 3E). The main effect of presentation condition and the functional localizer results were consistent with the findings of previous neuroimaging studies on dynamic facial expression processing, which reported heightened activity in the right pSTS and IFG, but not in the IPL (Liu et al., 2021; Zinchenko et al., 2018), as well as effective connectivity between the right pSTS and IFG (Sato et al., 2012, 2010). Similar results were also reported in a functional near-infrared spectroscopy hyperscanning study that compared the effects of dyad eye-to-eye (live eye contact) and eye-to-picture gaze (eye contact with the eyes in a picture) between two participants. Live eye-to-eye contact evoked greater responses in the left IFG pars opercularis and pre- and supplementary motor cortices. Psychophysiological interaction analysis showed greater functional connectivity between the left IFG and left superior temporal gyrus. The authors postulated that eye gaze alone evokes less face recognition, emotion processing, and social inferencing than dynamic facial expressions; therefore, they obrained left-dominant results (Hirsch et al., 2017). The parametric modulatory effect of mimicking EMG activity in the MNS in the present study was also observed in a study of simultaneous fMRI-EMG acquisition that investigated dynamic vs. static facial expression processing; the positive correlation between ZM and fMRI ROI activity for the happy-dynamic condition was significant in the right pSTS and showed a trend in the right IFG, whereas negative correlation between CS and ROI activity for the happy dynamic condition was significant in both the left pSTS and left IFG. In the anger-dynamic condition, there was a positive trend between CS and ROI activity for the right IFG (Rymarczyk et al., 2018). Anatomical evidence also showed that the IFG and pSTS are connected directly via the arcuate fasciculus and indirectly via other regions including the IPL and posterior temporal lobe (Friederici, 2009). These results were also consistent with the hypothesis that, the pSTS-IFG pathway in the MNS is involved in mimicry behaviors (Hamilton, 2008), and also consistent with findings showing that in macaques, visual input consisting of facial communicative signals bypasses the MNS node of the IPL (Ferrari et al., 2017), which is the indirect goal-emulation and action planning pathway (Hamilton, 2008). Together with these data, our results showing heightened activity in the right pSTS, IFG, and premotor cortex; parametric correlation with mimicking amplitude involving these regions; and modulated effective connectivity between these regions in the live condition indicate that, during the perception of live facial expressions, the MNS in the right hemisphere became more engaged in mirroring or mimicry processing than occurred with the observation of pre-recorded videos of other individuals.

The live effect has also been described as the audience effect, referring to a change in behavior caused by the belief that one is under observation (Hamilton and Lind, 2016). Alternatively, the watching eyes effect refers to the attention capture, self-referential processing and prosocial behaviors elicited by the perception of direct gaze (Conty et al., 2016). These behavioral effects have been associated with the metacognition in social interaction for the purpose of reputation management (Frith, 2012; Frith and Frith, 2011; Tennie et al., 2010). Zajonc also used drive theory to explain how the presence of conspecifics increased individuals’ arousal and influenced their performance of tasks (Zajonc, 1965). Considering the multitude of cognitive affective processes evoked by the audience and watching eyes effects, it is equally plausible that the live conditions examined in this study appeared to be more socially salient for the participants, increasing the subjective experiential arousal, as reflected in the significant presentation condition effect in arousal ratings and the enhanced salience network activity in the right amygdala in the [live > video] contrast (de Gelder et al., 2012; Jacobs et al., 2012; Leitão et al., 2021; Pourtois et al., 2013). Furthermore, modulated effective connectivity also corresponded to attention or executive control in the frontoparietal network (Fig. 3E), which was enhanced by heightened social cognition, especially the frontal eye field (BA 8) included in the IFG cluster and inferior parietal sulcus (Corbetta, 1998; Decety and Sommerville, 2003; Ptak, 2012; Sato et al., 2009; Scolari et al., 2015). The left pSTS, which was also more active in the live condition than in the video condition (Table 5, Fig. 3A), has been reported to modulate social attention for self-relevance (Sui et al., 2013). The univariate and connectivity results for the MNS and amygdala correspond to the taxonomically relevant yet parallel and undifferentiable cognitive affective processes elicited by the live effect beyond mirroring and mimicking.

### 4.3 Live effects modulated the mentalizing network in multivariate activation patterns

Although the mentalizing network, including the bilateral mPFC/ACC and PCC (Fig. S3C), which is involved in dynamic facial expression processing as identified by the functional localizer task, showed no difference between different presentation conditions in the passive observation task according to univariate analysis, the multivoxel activation patterns in these regions were distinguished between presentation conditions by the SVM algorithm, albeit with a small effect size (AUC). Whole-brain MKL analysis further indicated that this differentiation was mainly due to the activation patterns in the right ACC of the mentalizing network, which can be explained by metacognitive “we-mode” activation (Frith, 2012), which is the automatic taking account of the knowledge and intentions of others, in live conditions, as well as the watching eyes effect (Conty et al., 2016), resulting in greater expression of mimicry and engagement in social reputation management for the purpose of increasing likeability and social rapport (Hess and Fischer, 2013). Mentalizing computations resulting from the belief of being watched have been reported to occur in the pSTS/middle temporal gyrus, mPFC, and orbitofrontal cortex. Self-referential processing resulted in self-awareness and pro-social behaviors involved medial cortical structures such as the ACC and dorsal mPFC (dmPFC) (Conty et al., 2016).

### 4.4 The valence effect: Positive vs. negative

In terms of the main effects of emotion condition, we found the right IFG to be more active in the positive than the negative condition (Table 5, Supplementary Fig. S2D). According to a previous meta-analysis (Lindquist et al., 2016), the IFG is valence-general, i.e., it responds to both positive and negative stimuli. In the present study, positive stimuli were generally rated as further from the neutral level and more arousing than negative stimuli, regardless of whether the performance was pre-recorded or live (Tables 1 and 2, Fig. 2A, B). It is plausible that the positive stimuli engaged more valence-associated processing in the right IFG, which is particularly associated with emotional dynamic facial expression processing, especially in the context of social interaction (Liu et al., 2021). Comparisons in both directions between positive and negative stimuli showed significant occipital involvement (Supplementary Fig. S2D). However, the significant cluster for the positive > negative contrast had a higher position (larger MNI z-axis value) than the cluster in the negative > positive contrast. As the visual salience for the positive stimulus (moving mouth) had a lower position of than those for the negative stimulus (moving eyebrows), and inverse retinotopy of the primary visual cortex, these results are likely to reflect the effects of low-level visual features in dynamic facial expressions. Our laterality analysis showed no lateralized activation related to the valence effect (Results 3.3.3, Table 6).

**Table 6.** fMRI GLM laterality analysis results

| Regions | Cluster size | p (FWE-corr)* | T | B.A. | [x, y, z] |
| --- | --- | --- | --- | --- | --- |
| **Live > Video, Right > Left** |  |  |  |  |  |
| Middle occipital gyrus | 119 | 2.4724e-07 | 5.93 | 19 | 30 -82 -13 |
| Lingual gyrus |  |  | 4.77 | 18 | 6 -91 -7 |
| Lingual gyrus |  |  | 4.42 | 17 | 3 -79 -13 |
| Middle occipital gyrus | 57 | 2.6198e-04 | 5.32 | 17 | 15 -94 14 |
| Cuneus |  |  | 4.74 | 18 | 6 -97 14 |
| Middle occipital gyrus |  |  | 4.03 | 18 | 27 -88 17 |
| Precentral gyrus | 40 | 0.0026 | 5.23 | 6 | 51 5 41 |
| Precentral gyrus |  |  | 4.71 | 6 | 48 -4 53 |
| Middle temporal gyrus | 84 | 1.0222e-05 | 5.12 | 37 | 54 -58 2 |
| Inferior temporal gyrus |  |  | 4.50 | 37 | 51 -67 -4 |
| Middle temporal gyrus |  |  | 3.68 | 37 | 42 -64 -1 |
| Middle temporal gyrus | 64 | 1.0837e-04 | 4.78 | 21 | 48 -25 -7 |
| Middle temporal gyrus |  |  | 4.71 | 22 | 60 -40 2 |
| Middle temporal gyrus |  |  | 4.36 | 21 | 51 -31 -1 |
| **Live > Video, Right < Left** |  |  |  |  |  |
| Lingual gyrus | 48 | 8.6113e-04 | 4.93 | 18 | 15 -73 -4 |
| Cuneus |  |  | 4.71 | 17 | 12 -79 2 |
| * Cluster-level FWE-corrected for the whole-brain, cluster-forming threshold: uncorrected $p = .001$.<br>** All peak coordinates reported were listed in the right hemisphere, because the analysis was a comparison between contrasts of non-flipped and flipped images limited to the right hemisphere. However, the results should be interpreted as difference between hemispheres.<br>*** $n = 44$<br>Abbreviations: IFG = Inferior frontal gyrus; $p$ = p-value; $T$ = T-value; B.A. = Brodmann area; $x, y, z$ = MNI coordinates |  |  |  |  |  |

### 4.5 The behavioral and neural basis of social alignment

Spontaneous mimicry, emotional contagion, emotional and cognitive empathy, and social conformity have been incorporated under the framework of social alignment for dyadic interaction and group herding (Shamay-Tsoory et al., 2019). Motor synchrony, emotional contagion, and cognitive synchrony reciprocally influence one another and contribute to social alignment. Under this model, the dmPFC and dACC (mentalizing system), and anterior insula, serve as error detectors for social alignment. When misalignment is detected, the human MNS (IFG, IPL, pSTS, and premotor cortex) functioning in observation-execution act to align the emotion and motor behavior with those of the other person(s). The aligned and synchronized motor, emotion, and cognitive status reduce the prediction error in social alignment and herding (Shamay-Tsoory et al., 2019). Our spontaneous facial mimicry (EMG) and emotional contagion (ratings) findings implied that there is greater social alignment effort under the second-person (live) condition. Our contrasting neuroimaging findings between live and pre-recorded conditions further implied that social alignment may engage the misalignment detection and alignment enhancement networks in different ways neurocognitively. Within the MNS for alignment enhancement, the response to live interaction is presented as enhanced univariate activation amplitude, which was correlated with mimicking muscular activity amplitudes (Table 5, Fig. 3C) and enhanced effective connectivity within the MNS (Fig. 3E). However, we did not observe response amplitude difference in the mentalizing network for misalignment detection. Instead, the differentiable co-activation patterns of voxels implied different functional organization for misalignment detection between live and pre-recorded conditions. Note that although the observation-execution process of the MNS is explicitly described as a consequence of alignment error detection in the abovementioned framework, mirroring activity can also provide an embodied copy (Wood et al., 2016) of the observed motor, emotion, or cognitive status for gap detection in the mentalizing network. Such iterative alignment processes have been proposed and computationally implemented in the Hierarchical Modular Selection and Identification for Control (HMOSAIC) model (Wolpert et al., 2003). In HMOSAIC, the observer’s internal social model infers the hidden mental state of the observed person by predicting possible actions and their motor commands in the MNS and motor system, and comparing the predicted outcome with the observed action, likely to occur in the mentalizing network, leading to updated predictions about the internal model of the observed person and possibly leading to the alignment of both the observer’s internal model and the simulated internal model regarding the observed person. Although our neuroimaging data did not directly address the computational nature of different networks, our findings corroborated previous findings that knowledge of the mere presence of another individual could enhance emotion and autonomic synchronization (Golland et al., 2015) for social alignment. The present study has also provided empirical neuroscientific evidence for the ecological validity of the second-person approach for social neuroscience.

During social alignment, emotional contagion changed the trajectory of an agent’s emotional state (Gross, 2015). As both motor mirroring and emotional empathy facilitate emotional contagion (Hatfield et al., 2014; McIntosh, 2006), the alignment process can involve emotion appraisal (Paulus et al., 2013) and constructive processes (Gross and Feldman Barrett, 2011), and the resulting emotional state would be empathic, vicarious, and simulated (Paulus et al., 2013). Indeed, although our affective ratings and EMG results showed interactions between emotion and presentation conditions, univariate activity and effective connectivity in the MNS and amygdala were modulated by the presentation condition. Our MKL multivariate results imply that the right ACC and right putamen contributed 79.9% of the whole-brain differentiable multivoxel pattern between presentation conditions. According to the appraisal-by-content model, the mentalizing network, particularly the dmPFC, may also contribute to appraising the mental status, intentionality, and traits of others (Dixon et al., 2017). As vividly demonstrated in a previous fMRI study, in participants who read short stories involving four protagonists, the multivoxel activation pattern in the dmPFC could differentiate between representations of personality types (Hassabis et al., 2014). Conversely, the putamen has been shown to be associated with negative emotion suppression (Vanderhasselt et al., 2013), and anger suppression for the purpose of cooperative behavior (Eimontaite et al., 2019). Bilateral putamen volume has also been reported to be negatively associated with the accurate recognition of fearful faces (Uono et al., 2017). This finding is consistent with the role of the mentalizing network in social alignment error detection and the role of the MNS in alignment enhancement.

### 4.6 Right-dominant live effects

This study was not designed to determine laterality in live social cognition, as we did not control for basic relevant factors such as participant handedness. However, because the significant live effects were mainly in the right hemisphere, we decided to perform additional laterality analysis on contrasts to test for laterality before discussing the observed right dominance. We found right dominant live effects in the premotor cortex (BA 6), posterior (BA 37) and middle (BA 21 and 22) temporal lobe, and visual cortex. In the whole-brain MKL, both regions that contributed most to the classification weight were also in the right hemisphere (right ACC and putamen).

The visual spatial frequency hypothesis proposed that in the right-handed population, the right hemisphere is more sensitive to low spatial frequency (holistic processing of faces and objects), whereas the left hemisphere is more specialized at high spatial frequencies (written word processing) (Brederoo et al., 2020; Cai et al., 2013; Gerrits et al., 2019; Hellige, 1996). The analogous asymmetric sampling in time hypothesis (Poeppel, 2003) proposed that the left hemisphere preferentially processes information in short temporal integration windows (∼20–40 ms, e.g., phonetics), and right hemisphere homologs preferentially process information in long integration windows (∼150–250 ms, e.g., prosody or music melodies) (Flinker et al., 2019; Giroud et al., 2020; Hamilton, 2019; Zatorre et al., 1992). Previous studies have attributed these phenomena to the developmental (Howard and Reggia, 2007) or interhemispheric anatomical asymmetry (Deoni et al., 2011; O’Muircheartaigh et al., 2013; Toga and Thompson, 2003). Regarding stimuli applied in the present study, the face is of lower spatial frequency (Goffaux and Rossion, 2006), and would be associated with right-dominant processing according to the spatial frequency hypothesis. Dynamic facial expressions require integration windows of hundreds of milliseconds (Jack et al., 2014), and would therefore be right-dominant in processing according to the asymmetric sampling in time hypothesis.

These lower-level principles could explain other hypotheses relevant to our findings. It has been proposed that social stimuli tend to evoke right-dominant activation (Amaral et al., 2015; Vrtička et al., 2013), and the live conditions, which evoke the audience and watching eyes effects, could engage more social cognition and social attention control. Another possible explanation is that emotion processing is right-dominant (Sackeim et al., 1978), and the live conditions evoked stronger affective arousal, and enhanced emotion processing in the right hemisphere. However, the findings of previous neuroimaging studies do not support this hypothesis (Lindquist et al., 2016, 2012; Wager et al., 2003). In both cases, social stimuli usually consisted of human face or body movements, and emotion signals often consisted of facial expressions, speech prosodies or music melodies, which would evoke right-lateralized processing according to low-level spatial or temporal frequency principles. It is also possible that through associative learning (Keysers and Gazzola, 2014), higher level social or emotion processing became generally right-lateralized, albeit with more inter-individual variation, which was further enhanced by the top-down live effect (Conty et al., 2016; Frith, 2012; Hamilton and Lind, 2016) during interaction. Future studies should apply an integrative approach to clarify these taxonomically incoherent yet mechanistically overlapping, hypotheses of hemisphere lateralization.

### 4.7 Limitations and future directions

There are concerns that the effect of presentation condition on spontaneous facial mimicry, emotional contagion, and MNS and mentalizing network activation patterns could be driven by enhanced kinematic intensity of live model performances. Therefore, we used OpenFace to test whether models performed with stronger kinematic intensity in live conditions. The results showed that models actually smiled more weakly in the positive condition and frowned more weakly in the negative condition (Section 2.5). It was plausible that despite their best effort to maintain consistent intensity, the models were affected by fatigue and adaptation, which reduced their kinematic intensity, because the models made 50 live positive and 50 live negative facial expressions to each participant, and we tested three participants sequentially per session. Thus, it is unlikely that the enhancement effects that we reported in association with live performance were driven by bottom-up the kinematic intensity.

This study had certain limitations. To avoid confounding factors, such as models producing more spontaneous facial expressions and improvising facial expressions in live trials, the models posed with easily reproducible facial expressions. Consequently, the live and videotaped performances could not be differentiated without instruction (Hsu et al., 2020), and we could more confidently attribute the observed effect to participant awareness of the live interactions. However, in real-life interactions, facial expressions are usually embedded in the narrative flow of a conversation and seldom used in isolation (Bavelas and Chovil, 2018; Fernández-Dols, 2017). Posed, stereotyped facial expressions can also be perceived as less natural than evoked facial expressions (Faso et al., 2015). A more naturalistic social context and interactive activities such as conversation or games, may help simulate real-life social interactions. Such designs could evoke more natural emotional facial expressions and facial mimicry in dyadic interactions.

TThe passive viewing task prompted many participants to close their eyes during the experiment. During eye closure, the BOLD activity of participants was decoupled from the study paradigm, which resulted in noise and error in univariate analyses, as well as reduced efficacy of training and validation in MVPA. Although we reviewed the video recording of participants in the scanner, it is difficult to differentiate sleep from prolonged blinking or the slit-like eye-opening when participants are trying hard to keep their eyes open. Hence, we considered it impractical to determine the wakefulness state of the participants. As a result, we excluded runs with prolonged eye-closed periods. This approach reduced the quantity of data with non-task-associated-BOLD signals, but also significantly reduced the statistical power for univariate analysis and quantity of data available for machine learning. We performed an alternative analysis with more stringent exclusion criteria in the supplementary material, which showed results largely consistent with those reported in the main text. Because we created different pseudorandomized presentation sequences and jitter intervals for the participants, we believe that the duration of eye closure time did not differ significantly between the live and video conditions. Furthermore, previous decoders for sleep using fMRI or EEG-fMRI data relied on functional connectivity patterns between parcellated brain areas (Altmann et al., 2016; Tagliazucchi and Laufs, 2014), not regional multivoxel patterns. Thus, it is less likely that voxel patterns, and even less likely that voxels in our ROIs (i.e., mPFC and PCC alone), reflect sleep patterns instead of task-associated patterns. Designs including oddball tasks or regular button press might counteract the tendency to fall asleep, but there are concerns that diverting the attention of participants to non-social stimuli and tasks would interfere with the social cognition required for the task. As mentioned previously, a more interactive task design could also help to overcome this issue.

Although the task-associated multivoxel patterns in the mPFC/ACC and PCC could be classified using the SVM algorithm, the classifier performance was not outstanding. Our results do not suggest the presence of a reliable neural signature of the “we-mode” (Frith, 2012) across participants in the mentalizing network. There might be individual differences in the discriminability and homogeneity of the state representations of live vs. video conditions, which could be modulated by personality traits such as trait empathy. The performance might also be affected by a block design, increased quantity of collected data, and more sophisticated classifier algorithms. Future studies should investigate individual differences (e.g., trait empathy) of neural representation of “we-mode”, which would require a substantially larger sample size than the present study (Dubois and Adolphs, 2016).

To avoid the effects of age and sex on spontaneous facial mimicry (Hess and Fischer, 2013; Seibt et al., 2015), we recruited only young, female participants. Future studies should investigate the effects of age, sex, race, social groups, and socioeconomic status in dyadic pairs to determine the generalizability of our findings.

## 5. Conclusion

Our results provided clear empirical evidence for the theoretical claims about the ecological validity of switching from a third-person approach to a second-person approach in social neuroscience studies (Redcay and Schilbach, 2019). Our findings demonstrated changes in activation dynamics and functional organization at both network and regional levels, implying that the second-person approach could bring additional neurocognitive insight for neuroimaging studies in social neuroscience.

## Acknowledgement

This project was generously supported by the Japan Science and Technology Agency CREST (JPMJCR17A5) and JST Mirai Program (JPMJMI20D7). The authors would like to thank Mami Fujikura, Yuko Kuroda, Kazusa Minemoto, Mei Nakabayashi, and Masaru Usami (in alphabetic order) for their assistance. This study was conducted using the MRI scanner and related facilities of the Institute for the Future of Human Society, Kyoto University.

## Author Contribution

C.-T. H., W.S. T.K. and S.Y. conceived the experiment. C.-T. H., W.S., K.A., N.A. and R.N. prepared the experimental setup. C.-T. H. conducted the experiments and analyzed the results. C.-T. H., W.S. T.K., K.A., N.A., R.N. and S.Y prepared the manuscript. W.S. and S.Y. acquired the funding.

## Declaration of Interest

The authors declare no competing financial or non-financial interests.

## Data and code availability

The following data tables and codes are available on the Open Science Framework (https://osf.io/vybwm): rating data table; EMG response tables containing extracted EMG response values, excluding runs in which participants fell asleep and trials with inaccurate live facial expression performances; and R codes for statistical analyses of rating and EMG data. The fMRI group-level analysis t-maps and masks used for ROI time-series extraction and MVPA are available in the NeuroVault (https://identifiers.org/neurovault.collection:12818) (Gorgolewski et al., 2015). The original MRI datasets generated during the present study are not publicly available because the ethical approval and informed consent did not include permission for participant neuroimaging data to be publicly deposited. Based on the latest regulation in Japan, informed consent from all participants and approval from the ethics committee which initially approved the research project (the ethics committee of the Unit for Advanced Studies of the Human Mind, Kyoto University) are mandatory for sharing the original MRI datasets with a third party. The requirements are also subject to future policy changes of the Japanese Government and the Japan Science and Technology Agency.

## Notes

https://osf.io/vybwm/

https://identifiers.org/neurovault.collection:12818

### Competing Interest Statement

The authors have declared no competing interest.

### Summary of Updates

Additional analysis in the supplementary material with reduced sample size using a more stringent criteria for data exclusion based on the eye-closure of participants; additional limitation on the discriminability and effect size of MVPA results; additional limitation on the frequent eye-closure in the current paradigm. Updated data and code availability statements.

