## Supplementary material for "Enhanced Mirror Neuron Network Activity and Effective Connectivity during Live Interaction Among Female Subjects"

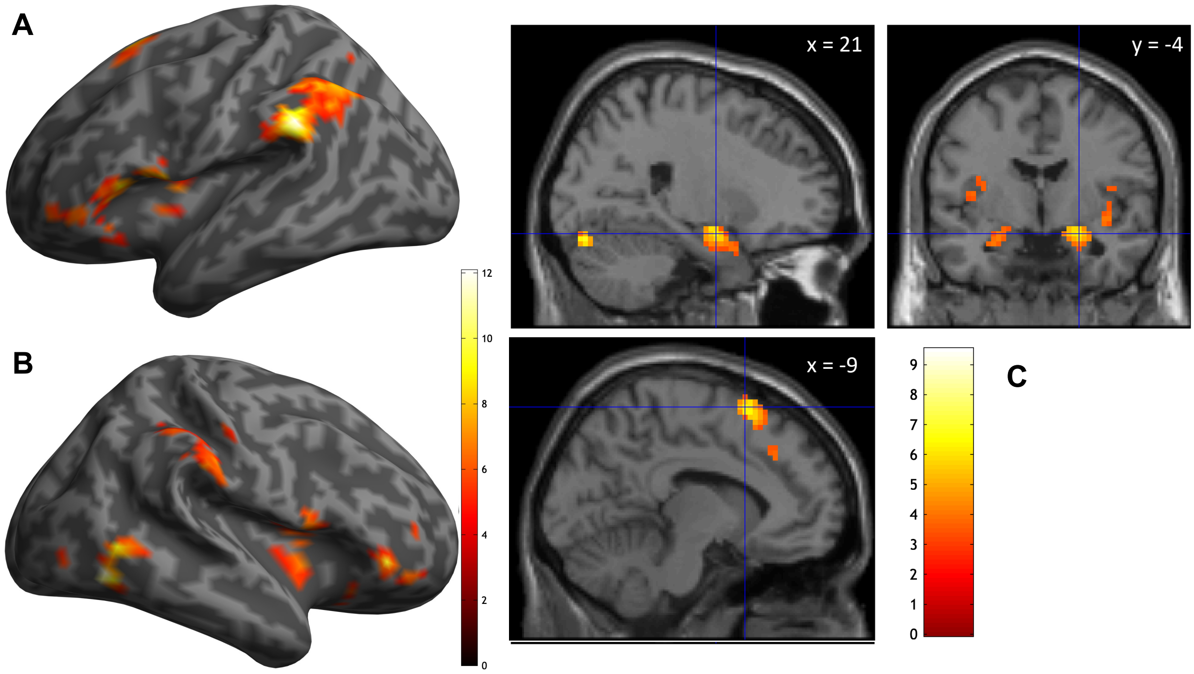


**Figure S1. Statistical parametric maps of average univariate activation in all four conditions in the passive viewing task.** (A) Lateral view of the left hemisphere showing activation of the left inferior frontal gyrus (IFG), supramarginal gyrus, and inferior parietal lobule (IPL). (B) Lateral view of the right hemisphere showing activation of the right IFG, insula, IPL, posterior temporal lobe, fusiform gyrus, and occipital pole. Colored bars indicate voxel *t* values. (C) Upper panels present section views with the crosshair at MNI [21 -4 -16] showing activation of bilateral amygdala. Left lower panel presents a sagittal section with the crosshair at MNI [-9 14 65] showing an activation cluster in the left supplementary motor area ($n = 44$).

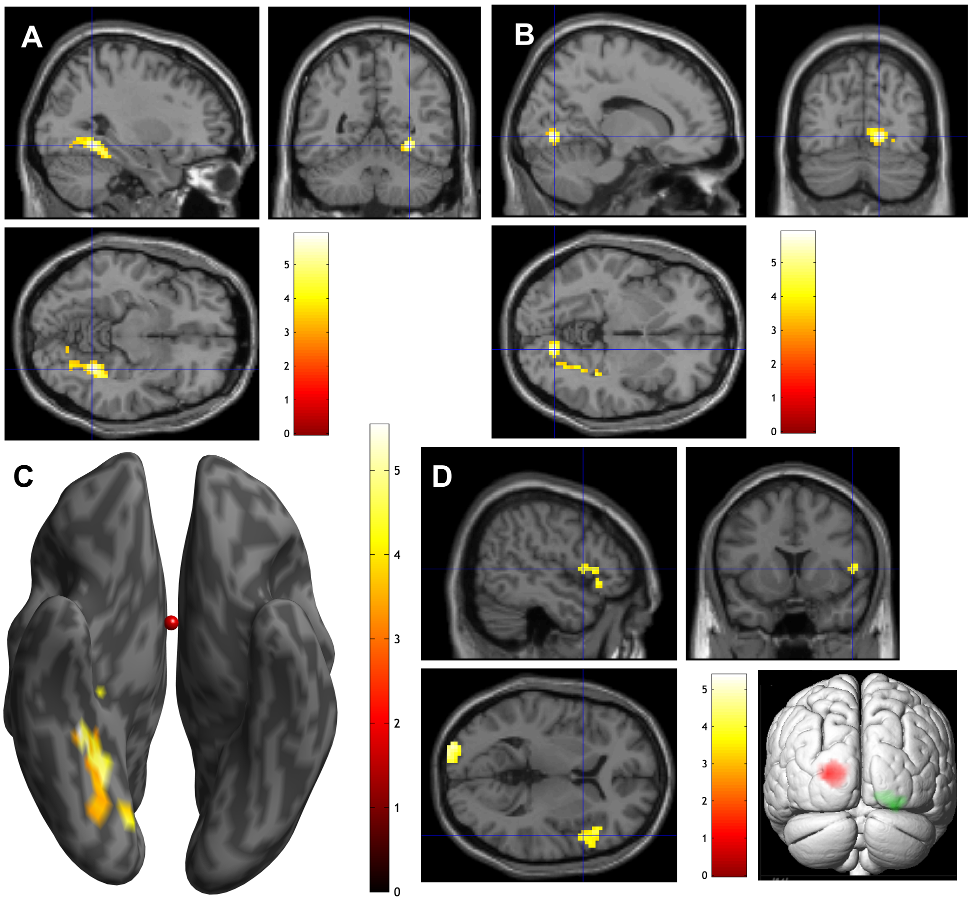


**Figure S2. Statistical parametric maps of passive observation task results.** (A-C) Main effects of presentation condition [Video > Live] in section views with crosshairs highlighting the right fusiform gyrus (panel A, MNI [30 -52 -10]) and right lingual gyrus (panel B, MNI [15 -73 -4]). (C) Bottom view shows two clusters of the contrast [Video > Live] (D) Main effect of emotion. Upper and left lower panels show the section view with crosshair, highlighting the IFG in the contrast [Positive > Negative] (MNI: [51 11 5]). Right lower panel shows posterior view of the clusters in the left upper occipital pole that were significant in the contrast [Positive > Negative] (red, peak MNI [-24 -100 8]) and a cluster in the right lower occipital pole that was significant in the contrast [Negative > Positive] (green, peak MNI [27 -88 -13]). Colored bars indicate voxel $t$ values ($n$ = 44).

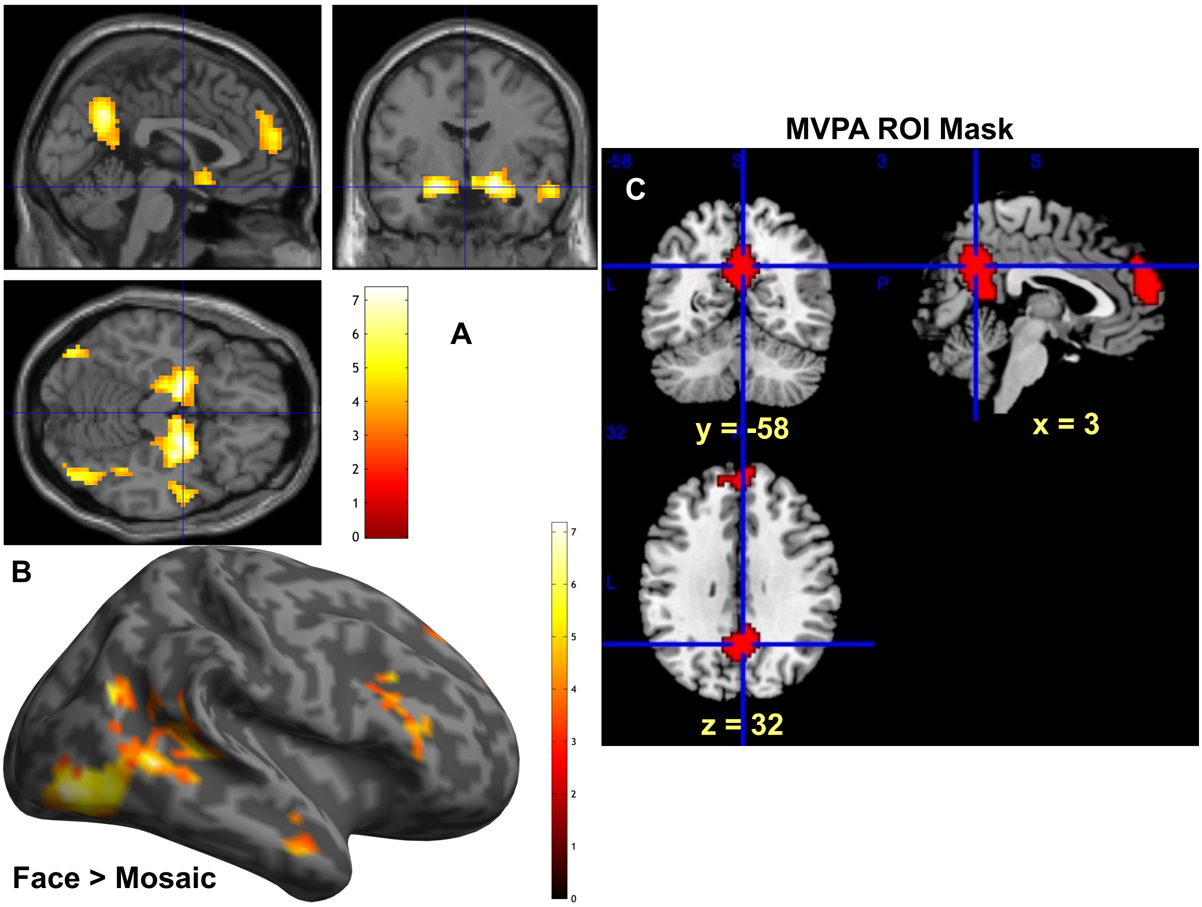


**Figure S3. Statistical parametric maps of localizer task contrast [dynamic faces > dynamic mosaics].** (A) Section views with crosshair at MNI [0 -4 -16] showing activation of the bilateral amygdala and hippocampus, bilateral fusiform gyrus, right anterior temporal lobe (aTL), dorsomedial prefrontal cortex (dmPFC), and posterior cingulate cortex (PCC). (B) Lateral view of the right hemisphere showing activation of the right IFG, aTL, fusiform gyrus, and dmPFC. Colored bars indicate voxel *t*-values ($n$ = 30). (C) Region of interest (ROI) mask of the mPFC/ACC and PCC used for the support vector machine (SVM) analysis. Upper left, upper right, and lower left panels show coronal, sagittal and transverse section views, respectively, with crosshair at MNI [3 -58 32].

**Table S1. fMRI GLM results of average activations in all four conditions.**

| H | Regions | Cluster size | p (FWE-corr)* | T | B.A. | [x, y, z] |
|---|---|---|---|---|---|---|
| **Beta > 0** | | | | | | |
| R | Middle occipital gyrus | 208 | 5.7346e-07 | 9.50 | 37 | 54 -67 -7 |
| | Middle temporal gyrus | | | 5.22 | 21 | 63 -58 5 |
| L | Inferior parietal lobule | 452 | 4.2911e-12 | 8.14 | 40 | -63 -43 32 |
| | Supramarginal gyrus | | | 5.68 | 39 | -45 -52 38 |
| | Inferior parietal lobule | | | 4.80 | 40 | -48 -46 53 |
| R | Middle frontal gyrus | 158 | 1.0598e-05 | 6.97 | 46 | 48 44 -4 |
| R | Fusiform gyrus | 60 | 0.0104 | 6.94 | 19 | 21 -85 -19 |
| L | Superior frontal gyrus | 158 | 1.0598e-05 | 6.45 | 6 | -9 14 65 |
| | Middle cingulate gyrus | | | 4.74 | 32 | -3 20 44 |
| | Medial frontal gyrus | | | 4.09 | 8 | -6 32 44 |
| R | Amygdala | 131 | 5.8177e-05 | 6.30 | | 24 -4 -16 |
| | Superior temporal pole | | | 4.58 | 28 | 27 8 -25 |
| | Posterior orbitofrontal cortex | | | 4.23 | 38 | 36 23 -16 |
| L | Amygdala | 51 | 0.0226 | 6.16 | | -24 -10 -16 |
| L | IFG pars opercularis | 364 | 2.1790e-10 | 6.04 | 44 | -57 11 5 |
| | IFG pars triangularis | | | 5.97 | 47 | -48 20 2 |
| | IFG pars opercularis | | | 5.39 | 44 | -51 11 11 |
| R | Insula | 213 | 4.3428e-07 | 5.42 | 13 | 42 8 -4 |
| | IFG, pars opercularis | | | 5.20 | 44 | 57 17 2 |
| | Insula | | | 4.98 | 13 | 36 2 11 |
| R | Inferior parietal lobule | 222 | 2.6473e-07 | 4.65 | 40 | 57 -37 50 |
| | Postcentral gyrus | | | 4.65 | 2 | 60 -22 32 |
| | Inferior parietal lobule | | | 4.40 | 40 | 51 -37 56 |

* Cluster-level FWE-corrected for the whole brain, cluster-forming threshold: uncorrected p = .001
** Multiple peaks were listed for one cluster
*** *n* = 44
Abbreviations: IFG = Inferior frontal gyrus; H = hemisphere; L = left; R = right; p = p-value; T = T-value; B.A. = Brodmann area; x, y, z = MNI coordinates

**Table S2. Localizer task results.**

| | | | | | | |
|---|---|---|---|---|---|---|
| **Face > Mosaic** | | | | | | |
| R | Amygdala & Hippocampus | 720 | $5.1292*10^{-14}$ | 7.35 | 20 | 21 -4 -16 |
| L | Hippocampus | | | 7.23 | 35 | -15 -7 -16 |
| R | Precuneus | 291 | $1.6667*10^{-7}$ | 7.08 | 23 | 3 -58 32 |
| R | Superior medial frontal gyrus | 249 | $9.9258*10^{-7}$ | 6.97 | 10 | 6 56 17 |
| L | Superior medial frontal gyrus | | | 3.95 | 32 | -9 53 29 |
| R | Inferior occipital gyrus | 677 | $1.9418*10^{-13}$ | 6.96 | 19/37 | 45 -79 -7 |
| L | Fusiform gyrus | 103 | 0.0014 | 6.86 | 19 | -39 -79 -13 |
| L | Cerebellum crus | 174 | $3.1676*10^{-5}$ | 6.81 | | -9 -82 -37 |
| R | IFG, pars triangularis | 101 | 0.0016 | 5.46 | 45 | 45 23 17 |
| R | aTL, middle temporal gyrus | 75 | 0.0078 | 5.41 | 21 | 60 -4 -19 |
| R | Superior medial frontal gyrus | 54 | 0.0325 | 4.84 | 9 | 9 44 41 |
| **Mosaic > Face** | | | | | | |
| L | Fusiform gyrus | 231 | $2.1981*10^{-6}$ | 8.28 | 19/37 | -27 -61 -13 |
| R | Fusiform & MOG | 919 | $1.1102*10^{-16}$ | 8.03 | 37/19/39 | 27 -49 -13 |

* Cluster-level FWE-corrected for the whole brain, cluster-forming threshold: uncorrected p = .001. Peak-level FWE-corrected for the small volume correction.
** Multiple peaks were listed for one cluster
*** $n = 30$
Abbreviations: aTL = anterior temporal lobe; IFG = Inferior frontal gyrus; H = hemisphere; L = left; MOG = middle occipital gyrus; R = right; p = p-value; T = T-value; B.A. = Brodmann area; x, y, z = MNI coordinates

**Supplementary Text S1**

Inclusion of task fMRI runs of participants with eyes closed for up to 50% of the time might be too liberal. Therefore, we performed additional analysis by including fMRI runs of participants with $< 25\%$ of time with their eyes closed. If $\geq 3$ runs were excluded, the participant was completely excluded from data analysis. This approach reduced the quantity of data with non-task-associated-blood oxygen level-dependent (BOLD) activity. However, it also substantially reduced the statistical power for univariate analysis and the quantity of data available for training and validation for MVPA. For the passive viewing task, data from 35 participants were analyzed. Of these participants, four runs for five participants and three runs for eleven participants were analyzed.

### S.1 Subjective Ratings

#### S.1.1 Valence Ratings

For the interaction effect of valence ratings, simple effects analysis showed that the positive-live condition (mean ± SE = 7.09 ± 0.1538, df = 35.1, 95% CI = [6.77, 7.40]) evoked greater positive emotions than the negative-live condition (mean ± SE = 3.36 ± 0.1083, df = 36.0, 95% CI = [3.14, 3.58]; difference = 3.72, SE = 0.236, df = 35.1, t = 15.785, $p < 0.0001$). Furthermore, the positive-video condition (mean ± SE = 6.05 ± 0.0939, df = 32.4, 95% CI = [5.86, 6.25]) evoked a greater positive emotion than the negative-video condition (mean ± SE = 3.78 ± 0.1206, df = 35.3, 95% CI = [3.53, 4.02]; difference = 2.28, SE = 0.172, df = 34.6, t = 13.244, $p < 0.0001$). Positive-live stimuli evoked stronger positive emotions than positive-video stimuli (estimate = 1.031, SE = 0.1231, df = 34.8, t = 8.371, $p < 0.0001$), whereas negative-live stimuli evoked stronger negative emotions than negative-video stimuli (estimate = -0.413, SE = 0.0913, df = 34.0, t = -4.525, $p = 0.0001$).

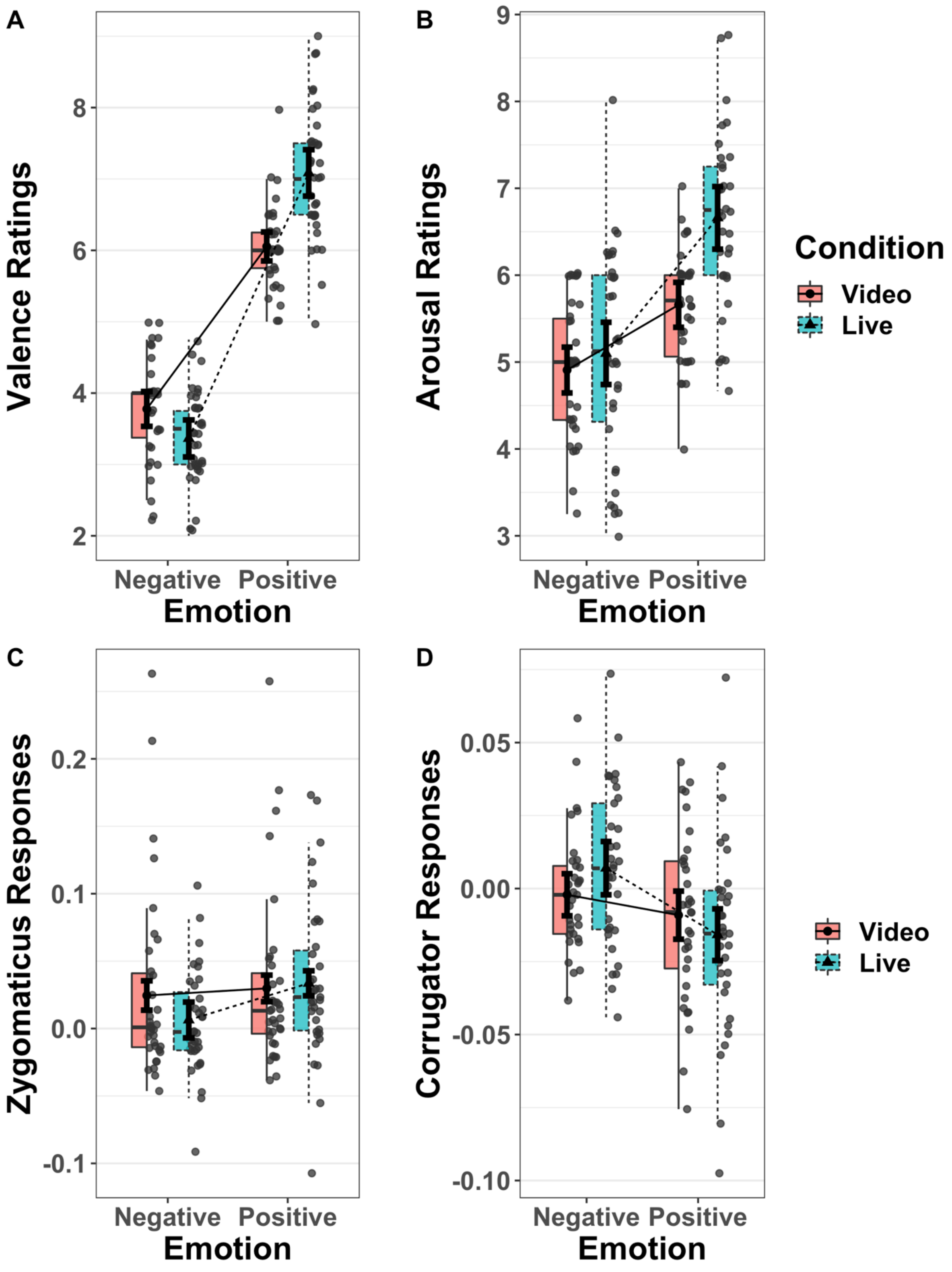


**Figure S4. Interaction between presentation and emotion conditions on ratings and facial muscular reactions.** (A) Valence ratings ($n$ = 35). (B) Arousal ratings ($n$ = 34). (C) Zygomaticus responses ($n$ = 35). (D) Corrugator responses ($n$ = 34). In all cases, significant interactions were observed between emotion conditions (positive vs. negative) and presentation conditions (video vs. live). For each condition, the right half shows the scattered dots of participant-wise mean values. Filled dots (video) and triangles (live) indicate group mean values.

Error bars indicate the within-subject standard error. The box on the left half indicates the median and the first and third quartiles of the distribution. The upper and lower whiskers extend from the hinge to the most extreme value, not exceeding 1.5 × the interquartile range (IQR) from the hinge.

**Table S3. Statistical summary of valence ratings with reduced sample size**

**Fixed Effects**

| Effect | Beta | 95% CI | SE | df | t-value | Pr(>\|t\|) | Rsq |
|---|---|---|---|---|---|---|---|
| Intercept | 5.0706 | (4.9638, 5.1778) | 0.0531 | 34.963 | 95.522 | < 2e-16* | |
| Emotion | 2.1209 | (1.8516, 2.3902) | 0.1337 | 35.075 | 15.868 | < 2e-16* | 0.878 |
| Presentation | 0.2185 | (0.1187, 0.3183) | 0.0495 | 35.359 | 4.412 | 9.19e-05* | 0.355 |
| Interaction | 0.7219 | (0.5555, 0.8891) | 0.0827 | 34.718 | 8.726 | 2.83e-10* | 0.687 |

**Random Effects**

| Group | Effect | Variance | SD | 95% CI | Corr. I. | 95% CI | Corr. E. | 95% CI | Corr. P. | 95% CI |
|---|---|---|---|---|---|---|---|---|---|---|
| Subject | Intercept | 0.0850 | 0.2916 | (0.2241, 0.3877) | | | | | | |
| | Emotion | 0.5979 | 0.7733 | (0.6146, 1.0057) | 0.08 | (-0.2783, 0.4194) | | | | |
| | Presentation | 0.0587 | 0.2423 | (0.1687, 0.3395) | -0.09 | (-0.4849, 0.3290) | 0.51 | (0.1225, 0.7804) | | |
| | E:P | 0.1852 | 0.4303 | (0.3165, 0.5868) | 0.49 | (0.1099, 0.7545) | 0.48 | (0.1226, 0.7332) | 0.42 | (-0.0284, 0.7459) |
| Residual | | 0.1882 | 0.4339 | (0.4043, 0.4671) | | | | | | |

*n* = 35, number of observations = 509

Formula: VAL ~ 1 + emotion_condition * presentation_condition + (1 + emotion_condition * presentation_condition | subject)

Abbreviations: CI: confidence interval; Corr. I.: correlation with the random effect of intercept; Corr. E.: correlation with the random effect of emotion condition; df: Satterthwaite approximations to degrees of freedom; Rsq: semi-partial $R^2$ effect size; SD: standard deviation; SE: standard error

### S.1.2 Arousal Ratings

For the interaction effect of arousal ratings, simple effects analysis showed that participants experienced greater arousal in response to the positive-live condition (mean ± SE = 6.66 ± 0.168, df = 34.0, 95% CI = [6.32, 7.00]) than the negative-live condition (mean ± SE = 5.10 ± 0.205, df = 34.0, 95% CI = [4.68, 5.52]; difference = 1.560, SE = 0.28, df = 34.1, t = 5.578, *p*

< 0.0001), as well as greater arousal in response to the positive-video condition (mean ± SE = 5.66 ± 0.106, df = 33.5, 95% CI = [5.44, 5.87]) than the negative-video condition (mean ± SE = 4.91 ± 0.131, df = 33.9, 95% CI = [4.64, 5.17]; difference = 0.751, SE = 0.19, df = 34.3, t = 3.957, $p$ = 0.0004). Participants also experienced greater arousal in response to positive-live stimuli than to positive-video stimuli (difference = 1.001, SE = 0.152, df = 33.7, t = 6.593, $p$ < 0.0001), whereas there was no difference in subjective arousal between negative-live and negative-video stimuli (difference = 0.191, SE = 0.156, df = 32.7, t = 1.223, $p$ = 0.2299).

**Table S4. Statistical summary of arousal ratings with reduced sample size**

**Fixed Effects**

| **Effect** | **Beta** | **95% CI** | **SE** | **df** | **t-value** | **Pr(>\|t\|)** | **Rsq** |
|---|---|---|---|---|---|---|---|
| Intercept | 5.5817 | (5.4094, 5.7544) | 0.0854 | 33.2934 | 65.330 | < 2e-16* | |
| Emotion | 0.8170 | (0.5117, 1.1220) | 0.1513 | 34.2185 | 5.399 | 5.11e-06* | 0.460 |
| Presentation | 0.4216 | (0.2621, 0.5811) | 0.0789 | 32.1678 | 5.343 | 7.20e-06* | 0.470 |
| Interaction | 0.4049 | (0.1905, 0.6202) | 0.1064 | 33.6075 | 3.806 | 0.000571* | 0.301 |

**Random Effects**

| **Group** | **Effect** | **Variance** | **SD** | **95% CI** | **Corr. I.** | **95% CI** | **Corr. E.** | **95% CI** | **Corr. P.** | **95% CI** |
|---|---|---|---|---|---|---|---|---|---|---|
| Subject | Intercept | 0.2257 | 0.4751 | (0.3692, 0.6286) | | | | | | |
| | Emotion | 0.7333 | 0.8563 | (0.6759, 1.1214) | -0.28 | (-0.5748, -0.0803) | | | | |
| | Presentation | 0.1670 | 0.4087 | (0.2992, 0.5605) | 0.66 | (0.3278, 0.8701) | 0.02 | (-0.3517, 0.3925) | | |
| | E:P | 0.2953 | 0.5434 | (0.3964, 0.7461) | -0.31 | (-0.6318, 0.0961) | 0.56 | (0.2053, 0.7731) | -0.03 | (-0.4405, 0.3834) |
| Residual | | 0.3005 | 0.5482 | (0.5097, 0.5917) | | | | | | |

$n$ = 34, number of observations = 483

Formula: ARO ~ 1 + emotion_condition * presentation_condition + (1 + emotion_condition * presentation_condition | subject)

Abbreviations: CI: confidence interval; Corr. I.: correlation with the random effect of intercept; Corr. E.: correlation with the random effect of emotion; df: Satterthwaite approximations to degrees of freedom; Rsq: semi-partial $R^2$ effect size; SD: standard deviation; SE: standard error

### S.2 Facial EMG

#### S.2.1 Zygomaticus major (ZM)

For the interaction effect of ZM responses, simple effects analysis showed that ZM reactions were stronger for the positive-live condition(mean ± SE = 0.03356 ± 0.0090, df = 36.4, 95% CI = [0.01513, 0.0518]) than the negative-live condition (mean ± SE = 0.00634 ± 0.0075, df = 37.1, 95% CI = [-0.00885, 0.0215]; difference = 0.02722, SE = 0.00658, df = 61.8, t = 4.138, *p* = 0.0001), whereas there was no difference in the ZM response between the positive-video (mean ± SE = 0.02976 ± 0.0114, df = 34.6, 95% CI = [0.00651, 0.0530]) and negative-video conditions (mean ± SE = 0.02449 ± 0.0101, df = 35.0, 95% CI = [0.00389, 0.0451]; difference = 0.00527, SE = 0.00654, df = 60.3, t = 0.805, *p* = 0.4238). The ZM contractions did not differ between positive-live and positive-video stimuli (difference = 0.0038, SE = 0.00645, df = 53.6, t = 0.589, *p* = 0.5582), whereas negative-live stimuli evoked greater ZM relaxations than negative-video stimuli (difference = -0.0182, SE = 0.00645, df = 53.7, t = -2.814, *p* = 0.0068).

**Table S5. Statistical summary of ZM reactions with reduced sample size**

**Fixed Effects**

| **Effect** | **Beta** | **95% CI** | **SE** | **df** | **t-value** | **Pr(>\|t\|)** | **Rsq** |
|---|---|---|---|---|---|---|---|
| Intercept | 2.354e-02 | (0.0062, 0.0410) | 8.631e-03 | 34.13 | 2.727 | 0.01002* | |
| Emotion | 1.149e-02 | (0.0035, 0.0196) | 3.978e-03 | 33.07 | 2.887 | 0.00680* | 0.201 |
| Presentation | -5.075e-03 | (-0.0131, 0.0027) | 3.889e-03 | 28.35 | -1.305 | 0.20239 | 0.057 |
| Interaction | 1.098e-02 | (0.0044, 0.0176) | 3.373e-03 | 5126 | 3.254 | 0.00114* | 0.002 |

**Random Effects**

| **Group** | **Effect** | **Variance** | **SD** | **SD 95% CI** | **Corr. I.** | **Corr. 95% CI** | **Corr. E.** | **Corr. 95% CI** |
|---|---|---|---|---|---|---|---|---|
| Subject | Intercept | 0.0025006 | 0.0500 | (0.0396, 0.0646) | | | | |
| | Emotion | 0.0003463 | 0.0186 | (0.0122, 0.0269) | 0.35 | (-0.0941, 0.6969) | | |
| | Presentation | 0.0003236 | 0.0180 | (0.0110, 0.0267) | -0.67 | (-0.9270, -0.2962) | -0.09 | (-0.5893, 0.3846) |
| Residual | | 0.0148254 | 0.1218 | (0.1194, 0.1242) | | | | |

*n* = 35, number of observations = 5223

Formula: ZM ~ 1 + emotion_condition * presentation_condition + (1 + emotion_condition +

presentation_condition | subject)

Abbreviations: CI: confidence interval; Corr. I.: correlation with the random effect of intercept; Corr. E.: correlation with the random effect of emotion; df: Satterthwaite approximations to degrees of freedom; Rsq: semi-partial $R^2$ effect size; SD: standard deviation; SE: standard error

#### S.2.2 Corrugator supercilii

For the interaction effect of CS responses, simple effects analysis showed greater CS responses in the negative-live condition (mean ± SE = 0.00699 ± 0.00461, df = 40.4, 95% CI = [-0.00232, 0.01631]) than in the positive-live condition (mean ± SE = -0.01580 ± 0.00529, df = 36.2, 95% CI = [-0.02652, -0.00508]; difference = 0.02280, SE = 0.00584, df = 51.1, $t$ = 3.902, $p$ = 0.0003), whereas no difference was detected in CS responses between the positive-video (mean ± SE = -0.00912 ± 0.00516, df = 37.0, 95% CI = [-0.01957, 0.00134]) and negative-video conditions (mean ± SE = -0.00212 ± 0.00365, df = 43.0, 95% CI = [-0.00949, 0.00525]; estimate = 0.00699, SE = 0.00585, df = 51.1, $t$ = 1.197, $p$ = 0.2370). There was no difference in CS responses between positive-live and positive-video conditions (difference = -0.00669, SE = 0.00446, df = 87.9, $t$ = 1.501, $p$ = 0.1371), whereas negative-live stimuli evoked greater CS contraction than negative-video stimuli (estimate = 0.00912, SE = 0.00456, df = 96.3, $t$ = -1.997, $p$ = 0.0486).

**Table S6. Statistical summary of CS reactions with reduced sample size**

**Fixed Effects**

| Effect | Beta | 95% CI | SE | df | t-value | Pr(>\|t\|) | Rsq |
|---|---|---|---|---|---|---|---|
| Intercept | -5.012e-03 | (-0.0183, 0.0045) | 3.274e-03 | 32.51 | -1.531 | 0.13545 | |
| Emotion | -1.053e-02 | (-0.0495, 0.0019) | 3.604e-03 | 29.59 | -2.922 | 0.00660* | 0.224 |
| Presentation | 8.590e-04 | (-0.0024, 0.0067) | 2.468e-03 | 33.09 | 0.348 | 0.72995 | 0.004 |
| Interaction | -7.901e-03 | (-0.0145, -0.0033) | 2.858e-03 | 5000 | -2.764 | 0.00572* | 0.002 |

**Random Effects**

| Group | Effect | Variance | SD | SD 95% CI | Corr. | Corr. 95% CI | Corr. | Corr. 95% CI |
|---|---|---|---|---|---|---|---|---|

| | | | | | I. | | E. | |
|---|---|---|---|---|---|---|---|---|
| Subject | Intercept | 2.830e-04 | 0.016823 | (0.0125, 0.0206) | | | | |
| | Emotion | 2.888e-04 | 0.016994 | (0.0068, 0.0245) | 0.43 | (-0.7196, 0.6298) | | |
| | Presentation | 6.419e-05 | 0.008012 | (0.0048, 0.0142) | 0.40 | (-0.2959, 0.4126) | -0.39 | (-0.3926, -0.3900) |
| Residual | | 1.037e-02 | 0.101822 | (0.0999, 0.1039) | | | | |

*n* = 33, number of observations = 5088

Formula: CS ~ 1 + emotion_condition * presentation_condition + (1 + emotion_condition + presentation_condition | subject)

Abbreviations: CI: confidence interval; Corr. I.: correlation with the random effect of intercept; Corr. E.: correlation with the random effect of emotion; df: Satterthwaite approximations to degrees of freedom; Rsq: effect size of semi-partial $R^2$ effect size; SD: standard deviation; SE: standard error

#### 3.3 fMRI Results

For the main effect of the presentation condition, the contrast [live > video] showed similar significant clusters to those of the results with 44 participants, including bilateral pSTS, right IFG and premotor cortex, and right amygdala. Additionally, the left IPL and cerebellum were significant clusters (Fig. S5A and B). The contrast [video > live] showed the same significant cluster in the right fusiform gyrus. For the main effect of the emotion condition, the contrast [negative > positive] showed the same cluster in the right occipital pole. There was no significant cluster that correlated with congruent EMG responses after sample size reduction.

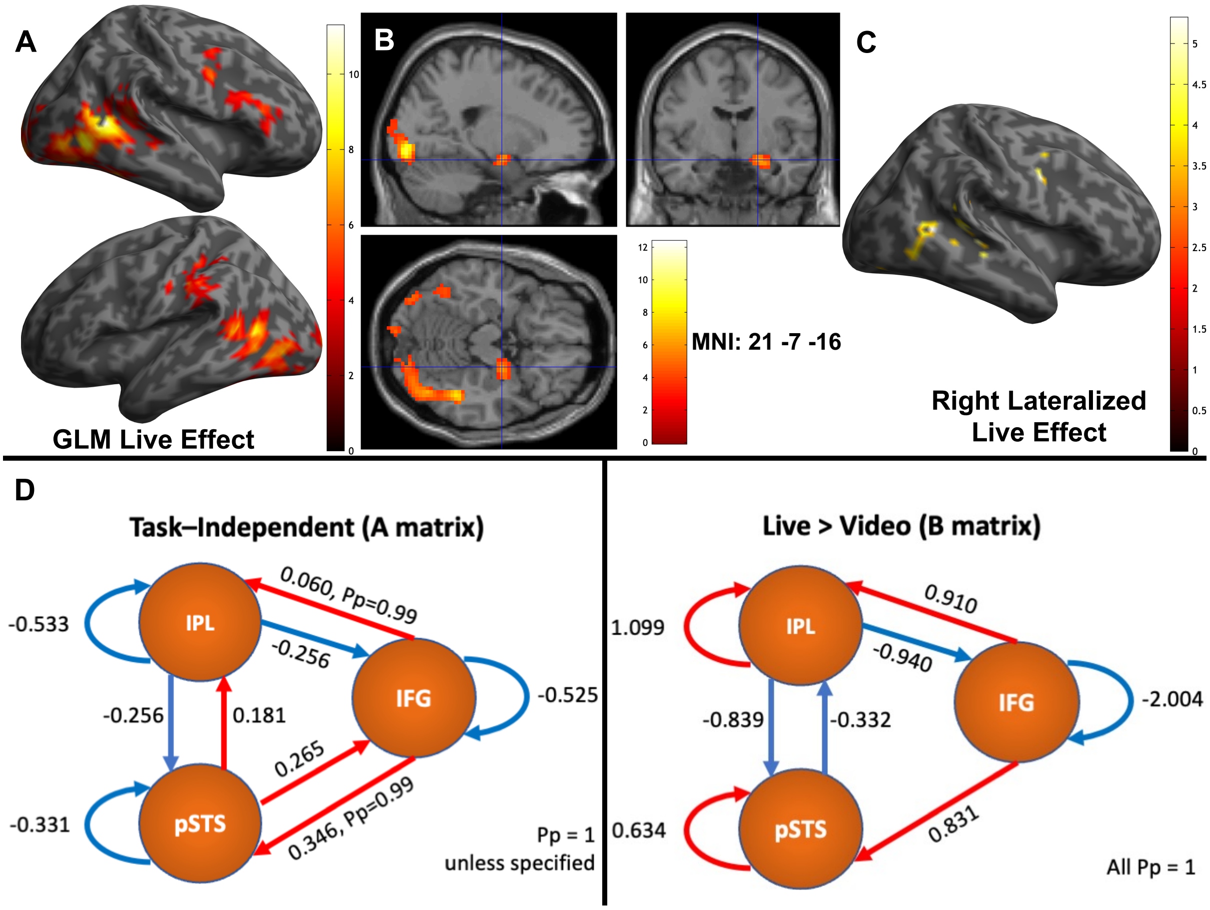


**Figure S5. Univariate fMRI and dynamic causal modeling (DCM) results of 35 participants.** (A) Statistical parametric maps of the main effects of presentation condition: [Live > Video]. Upper panel: lateral view of the right hemisphere showing activations in the right inferior frontal gyrus (IFG) and posterior superior temporal sulcus (pSTS). Lower panel: lateral view of the left hemisphere showing activation in the left pSTS (*n* = 35). (B) Section views with the crosshair (MNI = [21 -7 -16]), highlighting the right amygdala in the main effect of the presentation condition [Live > Video]. (C) Statistical parametric maps of significant right dominant clusters for the main effect of presentation condition: [Live > Video], with right lateral view of clusters in the right mid- and pSTS, premotor area, precentral gyrus and visual cortex. (D) DCM results of right MNS – IFG, pSTS, and inferior parietal lobule (IPL). The circles indicate three regions of interest (ROIs). Red and blue lines indicate positive and negative effective connectivity, respectively. Numbers indicate connectivity strength (in Hz). Left panel: task-independent baseline connectivity, estimated in the DCM as the A matrix. Right panel: Task modulatory effective connectivity, estimated in the DCM as the B matrix. In the design matrix, the parametric modulatory effect of live and video conditions were dummy-coded as 1 and -1, respectively (*n* = 35). Pp: posterior probability of parameter > 0.

### Table S7. fMRI GLM results with reduced sample size

| H | Regions | Cluster size | p (FWE-corr)* | T | B.A. | [x, y, z] |
|---|---|---|---|---|---|---|
| **Live > Video** | | | | | | |

| | | | | | | |
|---|---|---|---|---|---|---|
| R | Posterior middle temporal gyrus | 1394 | <1e-40 | 12.33 | 37 | 51 -58 8 |
| | Lingual gyrus | | | 10.87 | 18 | 18 -88 -7 |
| | Posterior middle temporal gyrus | | | 9.78 | 21 | 60 -52 5 |
| L | Posterior middle temporal gyrus | 590 | 2.5424e-14 | 8.89 | 39 | -54 -61 8 |
| | Middle occipital gyrus | | | 7.09 | 19 | -45 -79 -1 |
| | Posterior middle temporal gyrus | | | 7.03 | 37 | -48 -67 -1 |
| L | Lingual gyrus | 182 | 3.2301e-06 | 7.09 | 18 | -9 -97 -10 |
| | Middle occipital gyrus | | | 4.89 | 19 | -21 -100 11 |
| R | Amygdala | 63 | 0.0091 | 6.39 | | 21 -7 -16 |
| L | Postcentral gyrus | 203 | 9.8875e-07 | 6.28 | 3 | -54 -25 35 |
| | Inferior parietal lobule | | | 5.41 | 40 | -36 -34 41 |
| | Supramarginal gyrus | | | 5.13 | 40 | -54 -40 32 |
| R | IFG, premotor cortex | 433 | 1.5280e-11 | 6.19 | 6 | 48 5 38 |
| | Middle frontal gyrus | | | 5.91 | 6 | 45 -1 53 |
| | IFG, pars triangularis | | | 5.13 | 45 | 45 23 20 |
| L | Cerebellum Uvula | 59 | 0.0126 | 6.01 | | -18 -79 -37 |
| | Cerebellum Uvula | | | 4.72 | | -12 -76 -43 |
| **Video > Live** | | | | | | |
| R | Fusiform gyrus | 156 | 1.4932e-05 | 6.19 | 37 | 30 -52 -10 |
| | Lingual gyrus | | | 5.68 | 18 | 15 -73 -4 |
| | Parahippocampal gyrus | | | 5.68 | 37 | 36 -43 -7 |
| **Negative > Positive** | | | | | | |
| R | Lingual gyrus | 130 | 1.5927e-04 | 6.20 | 18 | 24 -88 -13 |

* Cluster-level FWE-corrected for the whole-brain, cluster-forming threshold: uncorrected p = .001.

** Multiple peaks were listed for one cluster if the cluster spans across both hemispheres.

*** $n = 35$

Abbreviations: IFG = Inferior frontal gyrus; MTG = middle temporal gyrus; STS = superior temporal sulcus; H = hemisphere; L = left; R = right; p = p-value; T = T-value; B.A. = Brodmann area; x, y, z = MNI coordinates

For the main effect of presentation conditions, the contrast [live > video] showed right-dominant activation [right > left] in the mid- and pSTS, premotor area, precentral gyrus and visual cortex (Fig. S5C), whereas a different subregion in the visual cortex showed left-dominant activation (Table S8), consistent with the results of the entire sample. There was no significant laterality for results of the main effect of emotion conditions, the interaction effect between two factors, or the parametric modulatory effect of facial mimicry.

**Table S8. fMRI GLM laterality analysis results with reduced sample size**

| Regions | Cluster size | p (FWE-corr)* | T | B.A. | [x, y, z] |
|---|---:|---:|---:|---:|---|
| **Live > Video, Right > Left** | | | | | |
| Middle occipital gyrus | 131 | 4.3415e-08 | 5.77 | 19 | 30 -82 -13 |
| Lingual gyrus | | | 5.18 | 18 | 18 -82 -7 |
| Lingual gyrus | | | 4.97 | 18 | 9 -91 -13 |
| MFG | 54 | 2.9137e-04 | 5.34 | 8 | 51 5 41 |
| Precentral gyrus | | | 5.29 | 6 | 48 -4 53 |
| MFG, premotor cortex | | | 4.86 | 6 | 45 5 50 |
| Superior temporal gyrus | 73 | 2.5518e-05 | 5.33 | 37 | 51 -55 8 |
| Inferior temporal gyrus | | | 4.41 | 37 | 51 -67 -4 |
| Middle temporal gyrus | | | 4.24 | 37 | 42 -64 -1 |
| Middle occipital gyrus | 34 | 0.0053 | 5.07 | 18 | 12 -94 11 |
| Superior temporal gyrus | 72 | 2.8842e-05 | 4.81 | 21 | 48 -28 2 |
| Middle temporal gyrus | | | 4.78 | 22 | 60 -40 2 |
| Superior temporal gyrus | | | 4.78 | 42 | 54 -37 11 |
| **Live > Video, Right < Left** | | | | | |
| Cuneus | 38 | | 5.11 | 18 | 18 -82 5 |
| Lingual gyrus | | | 4.60 | 18 | 15 -73 -4 |

* Cluster-level FWE-corrected for the whole-brain, cluster-forming threshold: uncorrected p = .001.

** All peak coordinates reported were listed in the right hemisphere, because the analysis was a comparison between contrasts of non-flipped and flipped images limited to the right hemisphere. However, the results should be interpreted as difference between hemispheres.

*** $n = 44$

Abbreviations: MFG = Middle frontal gyrus; p = p-value; T = T-value; B.A. = Brodmann area; x, y, z = MNI coordinates

ROIs in the DCM of reduced sample size were defined using the functional masks of the right

IFG and pSTS from the [live > video] contrast, and the right IPL from the average activation of four conditions of 35 participants. Group DCM results (Fig. S5D) showed that baseline effective connectivity consisted of positive effective connectivity between all pairs of nodes except for the negative effective connectivity from the right IPL to the right pSTS and from the right IPL to the right IFG. The modulatory effect of live over video conditions consisted of the positive effective connectivity from the right IFG to the right pSTS (0.831 Hz, Pp = 1), positive effective connectivity from the right IFG to the right IPL (0.910 Hz, Pp = 1), negative effective connectivity from the right IPL to the right IFG (-0.940 Hz, Pp = 1), negative effective connectivity from the right IPL to the right pSTS (-0.940 Hz, Pp = 1),and negative effective connectivity from the right pSTS to the right IPL (-0.332 Hz, Pp = 1).

Using the criteria of maximum eye closure time of 25%, only 24 participants remained in the localizer analysis. The contrast [dynamic face > dynamic mosaic] was significant in bilateral amygdala, right posterior temporal cortex, right IFG pars triangularis, and the mentalizing network including the mPFC and PCC/precuneus (Table S9). A functional mask containing the significant clusters of the right mPFC and PCC/precuneus was created for MVPA with the reduced sample size.

**Table S9. Localizer task results with the reduced sample size.**

| H | Regions | Cluster size | p (FWE-corr)* | T | B.A. | [x, y, z] |
|---|---|---|---|---|---|---|
| **Face > Mosaic** | | | | | | |
| L | Amygdala | 136 | 1.7761e-04 | 7.15 | | -18 -7 -16 |
| | Hippocampus | | | 6.03 | | -24 -16 -13 |
| | IFG, pars orbitalis | | | 5.07 | 11 | -27 11 -22 |
| R | Inferior occipital gyrus | 431 | 4.4427e-10 | 7.03 | 19 | 48 -76 -10 |
| | Superior temporal gyrus | | | 5.86 | 22 | 60 -55 8 |
| | Superior temporal gyrus | | | 5.24 | 21 | 51 -40 8 |
| R | Hippocampus | 148 | 9.3079e-05 | 6.98 | | 30 -10 -16 |
| | Amygdala | | | 6.78 | | 18 -4 -16 |

| | | | | | | |
|---|---|---|---|---|---|---|
| | Amygdala | | | 6.34 | | 24 2 -19 |
| R | Precuneus | 155 | 6.4366e-05 | 6.48 | 7 | 3 -58 29 |
| | Posterior cingulate cortex | | | 4.81 | 29 | 3 -49 14 |
| L | Inferior occipital gyrus | 75 | 0.0068 | 6.47 | 19 | -42 -82 -7 |
| R | IFG, pars triangularis | 50 | 0.0395 | 6.00 | | 36 17 23 |
| | IFG, pars triangularis | | | 5.49 | 45 | 45 23 17 |
| | IFG, pars triangularis | | | 5.06 | 45 | 51 32 14 |
| L | Cerebellum Pyramis | 55 | 0.0273 | 5.68 | | -9 -85 -37 |
| | Cerebellum Uvala | | | 4.47 | | -21 -79 -37 |
| R | Superior medial frontal gyrus | 93 | 0.0021 | 4.88 | 9 | 6 56 20 |
| | Superior medial frontal gyrus | | | 4.77 | 10 | 6 59 32 |
| | Superior medial frontal gyrus | | | 4.60 | 10 | 9 59 11 |
| **Mosaic > Face** | | | | | | |
| L | Fusiform gyrus | 146 | | 7.83 | 37 | -30 -46 -13 |
| | Lingual gyrus | | | 7.38 | 19 | -27 -61 -10 |
| R | Fusiform gyrus | 211 | | 7.34 | 37 | 27 -52 -13 |
| | Fusiform gyrus | | | 6.68 | 19 | 30 -67 -7 |
| R | Fusiform gyrus | 102 | | 6.11 | 19 | 39 -73 20 |
| | Middle occipital gyrus | | | 4.00 | 19 | 33 -85 29 |
| L | Posterior middle temporal gyrus | 112 | | 5.47 | 19 | -36 -82 17 |
| | Middle occipital gyrus | | | 5.18 | 19 | -27 -85 26 |
| R | Cuneus | 108 | | 4.69 | 18 | 15 -91 2 |
| | Cuneus | | | 4.51 | 19 | 12 -94 20 |
| | Cuneus | | | 4.08 | 18 | 18 -88 26 |

* Cluster-level FWE-corrected for the whole brain, cluster-forming threshold: uncorrected p = .001. Peak-level FWE-corrected for the small volume correction.
** Multiple peaks were listed for one cluster
*** *n* = 24
Abbreviations: IFG = Inferior frontal gyrus; H = hemisphere; L = left; R = right; p = p-value; T = T-value; B.A. = Brodmann area; x, y, z = MNI coordinates

With 35 participants, the binary SVM algorithm limited to the functional mask of right mPFC and PCC/precuneus yielded a group-level classification accuracy of 53.93% (area under the receiver operating characteristic curve (AUC) = 0.54, positive predictive values for classes [live, video] = [55.47%, 53.07%]). A permutation test, with 10,000 repetitions, showed that the accuracy was significantly higher than the chance level of 50% ($p = 2.9997 \times 10^{-4}$). The whole-brain MKL yielded a whole-brain group-level classification accuracy of 50% at the chance level.
